# Colitis-primed circulating T cells drive small intestinal barrier function through Nod2, microbiota and myosin light chain kinase-dependent mechanisms

**DOI:** 10.1101/2025.07.10.663215

**Authors:** Arthur Mauduit, Talal Nabhani, Luisa Morelli, Floriane Cherrier, Rosemary Abdoul Ahad, Nidal Nabhani, Gurminder Singh, Maryline Roy, Gilles Dietrich, Jerrold R. Turner, Jean-Pierre Hugot, Andrew Macpherson, Emmanuel Mas, Frédérick Barreau, Ziad Al Nabhani

## Abstract

Inflammatory bowel diseases (IBD), including Crohn’s disease (CD) and ulcerative colitis (UC), are characterized by intestinal inflammation and barrier dysfunction. While disruption of the intestinal barrier contributes to the pathogenesis of IBD, yet how colonic inflammation alters small intestinal homeostasis remains poorly defined. Here, we demonstrate that three models of colitis, including 2,4,6-Trinitrobenzenesulfonic acid (TNBS), Dextran Sulfate Sodium (DSS), and oxazolone, induce paracellular barrier dysfunction in the small intestine, with model-specific immune profiles. Th1/Th17-skewed responses (TNBS and DSS) were associated with microbiota-dependent upregulation of myosin light chain kinase (MLCK), RORγt⁺ CD4⁺ T cell expansion, and elevated expression of Nod2, which was not observed under Th2-dominant (oxazolone) conditions. Inhibition of MLCK restored barrier function and suppressed inflammation in a CD4⁺ T cell–dependent manner. Using Nod2-deficient and 2939insC mutant mice, we showed Nod2 as a critical regulator of small intestinal permeability during colitis. Bone marrow chimeras revealed compartment-specific roles of Nod2, with non-hematopoietic Nod2 sufficient to preserve epithelial integrity, while hematopoietic Nod2 expression is required to limit cytokine-mediated inflammation. Moreover, MLCK inhibition ameliorated intestinal lesions only in mice with Nod2 deficiency in the hematopoietic compartment. Finally, microbiota transfer experiments ruled out a causal role for dysbiosis in driving small intestinal permeability defects in Nod2-deficient mice. These findings uncover a Nod2–MLCK–CD4⁺ T cell axis linking colonic inflammation to small intestinal barrier dysfunction and highlight distinct immune–epithelial–microbial mechanisms shaping intestinal homeostasis during colitis.

**Graphical Abstract:** Circulating RORγt⁺ Foxp3⁻ CD4⁺ T cells from mice with TNBS- or DSS-induced colitis increase small intestinal permeability by promoting cytokine secretion that enhances MLCK activity. This effect is attenuated by Nod2 activation. Specifically, Nod2 expression in hematopoietic cells reduces cytokine secretion and permeability, while activation of Nod2 in non-hematopoietic cells alone is sufficient to prevent increased permeability by directly regulating MLCK activity, independently of cytokine influence. Moreover, the absence of Nod2 is associated with inflammatory lesions in the small intestine of TNBS-treated mice.

## Introduction

Crohn’s disease (CD) and ulcerative colitis (UC), the principal forms of inflammatory bowel disease (IBD), are chronic immune-mediated disorders of the gastrointestinal tract characterized by spatially distinct patterns of inflammation and divergent immunopathology. While CD typically involves transmural inflammation of the terminal ileum and colon and is associated with Th1/Th17 cytokine responses, UC primarily affects the colonic mucosa with a Th2-skewed immune profile. Although the colon is the principal site of inflammation in many experimental colitis models, increasing evidence indicates that intestinal inflammation can induce physiological alterations in remote sites, including the small intestine. Impaired small intestinal barrier integrity is increasingly recognized as a pathogenic factor in CD, contributing to disease recurrence, malabsorption, and microbial translocation (Turner, 2009). Previous work indicates that IBD patients, in remission or in relapses phases, have an elevated intestinal permeability. This increased permeability has been described in inflamed but also in non-inflamed mucosal area (Zeissig et al., 2007). However, the mechanisms linking localized colonic inflammation to remote small intestinal dysfunction remain largely undefined.

Experimental models of colitis have been instrumental in defining mucosal immune pathways in IBD. Among these, TNBS (2,4,6-trinitrobenzene sulfonic acid) and DSS (dextran sulfate sodium) elicit robust T helper 1 (Th1)/Th17 responses and mimic features of CD, whereas oxazolone induces a Th2-skewed inflammation reminiscent of UC. Whether these divergent immune responses differentially modulate small intestinal barrier function is unclear. While previous studies have largely focused on TNF-α and IFN-γ as key cytokines driving barrier defects (Al Nabhani et al., 2017b; Clayburgh et al., 2005; Su et al., 2009; Wang et al., 2005), other inflammatory mediators, including IL-1β, IL-12, and Th2 cytokines such as IL-4, IL-5, and IL-13, have also been implicated in regulating tight junction integrity, yet their roles in small intestinal barrier regulation during colitis are incompletely defined.

Tight junction remodeling in response to inflammation is known to involve the myosin light chain kinase (MLCK) pathway, which is activated by various cytokines and microbial stimuli. MLCK-dependent contraction of the peri-junctional actomyosin ring increases paracellular permeability and has been implicated in IBD pathogenesis (Blair et al., 2006; Graham et al., 2019). In fact, MLCK’s expression and activity in ileum and colon are increased in the areas affected by IBD. A positive correlation between this increase and active inflammation has been shown (Blair et al., 2006). Importantly, both microbial and immune-derived signals can regulate MLCK expression and activity, positioning it as a critical mediator of inflammation-induced barrier dysfunction. However, how immune polarization (Th1/Th17 vs. Th2), microbiota, and cytokine diversity converge on MLCK-mediated permeability in the small intestine remains unclear.

Among genetic factors linked to CD susceptibility (Podolsky, 2002), nucleotide-binding oligomerization domain-containing protein 2 (NOD2) plays a central role. NOD2 is highly expressed in both hematopoietic and non-hematopoietic compartments of the intestine and is critical for sensing bacterial peptidoglycans and maintaining epithelial barrier integrity. Mutations in *NOD2*, particularly the CD-associated 3020insC (1007fs) variant, are strongly associated with ileal CD and have been implicated in dysregulated immune responses, defective bacterial clearance, and impaired regulation of epithelial permeability. We previously shown that NOD2 expression is increased away from primary inflammatory lesions in naive pediatric CD patients and in the small intestine of mice with TNBS-induced colitis(Al Nabhani et al., 2020). Yet, the compartment-specific role of Nod2 in coordinating small intestinal barrier function during distal colonic inflammation is not fully understood.

TNBS-induced colitis is a well-known model of self-limited inflammation, studies have shown that rectal administration of TNBS to rats or mice can induce some limited changes in the small intestine but without overt histological lesions (Amit-Romach et al., 2006; Barreau et al., 2007). For instance, the villous surface area and the density of goblet-cells in colitis-induced rats was smaller in small intestine compared to control (Amit-Romach et al., 2006). In wild-type (WT) mice, we have recently reported an increased expression of pro-inflammatory cytokines in the small intestine with no overt duodenitis or ileitis (Al Nabhani et al., 2020). Moreover, T cell recirculation between the colon and small intestine represents an emerging mechanism by which distal inflammation may perturb remote epithelial function. CD4⁺ T cells, particularly RORγt⁺ Th17 cells, are known to expand in colitis and contribute to cytokine-driven barrier disruption (Mickael et al., 2022). Whether distinct T helper subsets exert differential effects on small intestinal permeability depending on the immune polarization of colitis remains to be elucidated.

In this study, we systematically dissect how different immune responses during colitis, Th1/Th17 (TNBS, DSS) versus Th2 (oxazolone), impact small intestinal homeostasis. We show that distinct cytokine environments promote MLCK-dependent barrier dysfunction in the small bowel via microbiota- and CD4⁺ T cell–dependent mechanisms. Furthermore, we identify Nod2 as a key regulator of small intestinal permeability and inflammation, acting through both hematopoietic and non-hematopoietic compartments. These findings highlight how immune polarization, microbial cues, and genetic susceptibility converge to shape intestinal barrier integrity beyond the site of primary inflammation.

## Results

### Distinct immune responses in colitis models differentially impact small intestinal homeostasis

To investigate how distinct immune responses during colonic inflammation influence small intestinal homeostasis, we evaluated the effects of three colitis models, TNBS, DSS, and oxazolone, on small intestinal integrity. These models elicit different immune profiles: TNBS and DSS induce a Th1-skewed response, characteristic of CD, whereas oxazolone promotes a Th2-polarized response, resembling that observed in UC. Adult C57BL/6J mice raised under Specific pathogen-free (SPF) conditions were co-housed for four weeks, then separated into groups and treated with TNBS or oxazolone via intra-rectal administration, or DSS through drinking water, to induce colitis. Control mice received corresponding vehicle treatments. As expected, induction of colitis in all three models resulted in robust inflammatory responses, evidenced by body weight loss, elevated disease activity index (DAI), shortened colon length, and increased macroscopic damage scores (*Fig. S1 A–D*). These pathological changes were accompanied by elevated expression of multiple inflammatory cytokines at the site of colonic inflammation (*Fig. S1 E–F*). In the duodenum and ileum, both mRNA and protein levels of inflammatory cytokines were significantly increased in TNBS-, DSS-, and oxazolone-treated mice compared to vehicle controls (*Fig. 1A–D*). Notably, TNF-α, IFN-γ, IL-1β, and IL-12 were upregulated across all colitis models (*Fig. 1A–B*), whereas Th2-associated cytokines, including IL-4, IL-5, and IL-13, were selectively elevated in the oxazolone-treated group (*Fig. 1C–D*). These cytokine elevations were associated with increased mRNA expression of *Tnfr2* (*Fig. 1E*) but not *Tnfr1* (*Fig. 1F*), and were linked to enhanced paracellular permeability, as indicated *in vivo* (*Fig. 1G*) and *ex vivo* using Ussing chamber analysis (*Fig. 1H*). Furthermore, expression of genes encoding tight junction proteins, such as *Tjp1* and *Tjp2*, was altered in duodenum and ileum of colitis-suffering mice (*Fig. 1I–J*). Together, these results demonstrate that all three colitis models disrupt small intestinal homeostasis by promoting inflammation and barrier dysfunction, with distinct cytokine signatures reflective of underlying Th1 or Th2 immune responses. These findings underscore the systemic impact of colonic inflammation on small intestinal physiology across different immune response types.

**Figure 1.**
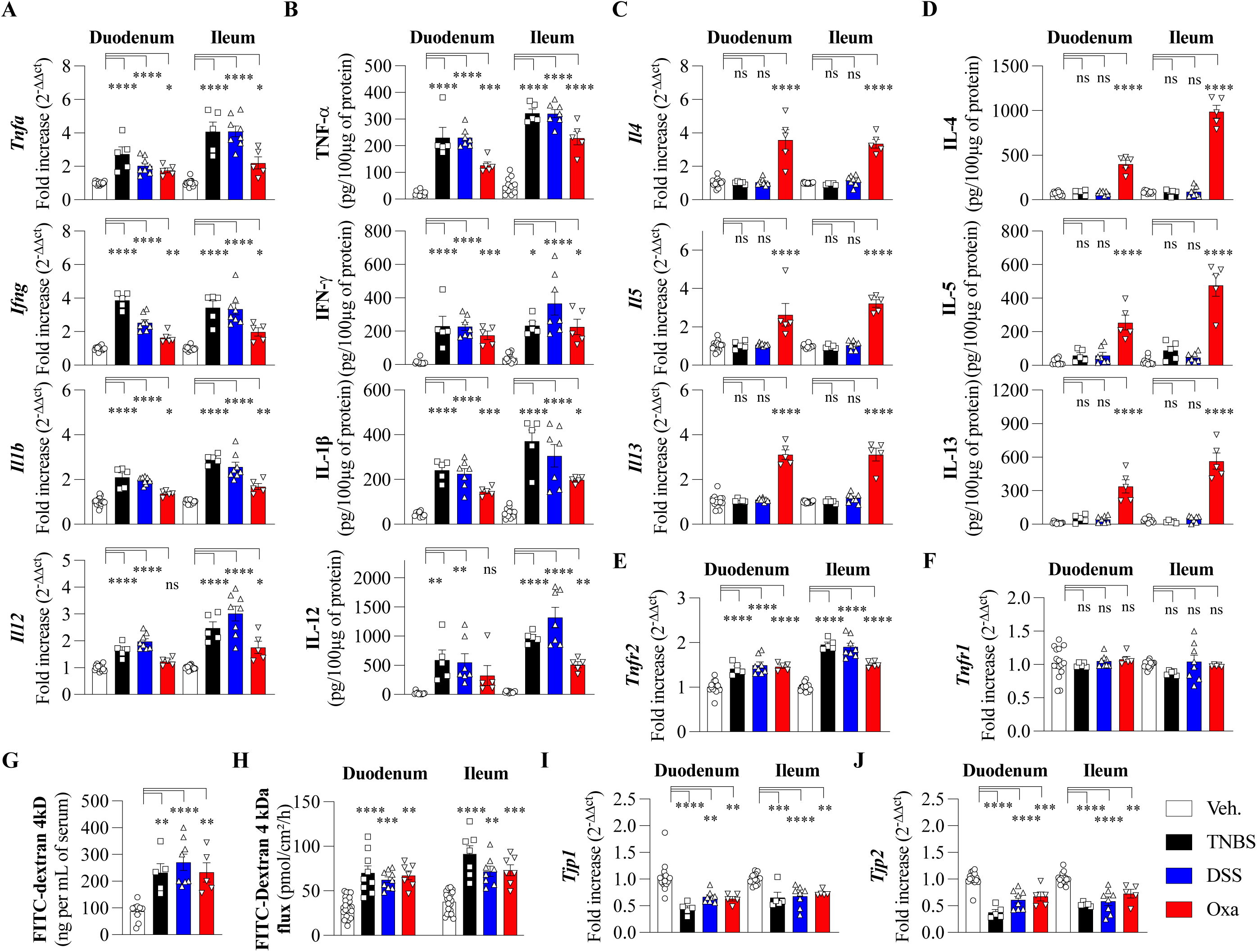
Colitis induced by TNBS, DSS, or Oxazolone alters cytokine expression and compromises duodenal and ileal barrier function. **(A-D)** Cytokine responses in the duodenum and ileum of WT mice with TNBS-(black), DSS-(blue), or oxazolone-induced (red) colitis. Control mice received vehicle only (white). **(A)** mRNA expression of *Tnfa*, *Ifng*, *Il1b*, and *Il12* measured by qPCR. **(B)** Protein levels of TNFα, IFNγ, IL-1β, and IL-12 measured by ELISA. **(C)** mRNA expression of *Il4*, *Il5*, and *Il13* measured by qPCR. **(D)** Protein levels of IL-4, IL-5, and IL-13 measured by ELISA. **(E and F)** mRNA expression of (E) *Tnfr2* and (F) *Tnfr1*. **(G)** Paracellular permeability measured by oral FITC-dextran gavage. **(H)** Duodenal and ileal paracellular permeability assessed by Ussing chamber. **(I and J)** mRNA expression of (I) *Tjp1* and (J) *Tjp2*. All data are from WT mice with colitis induced by TNBS, DSS, or oxazolone; controls received vehicle only. Each data point represents one mouse; *n* ≥ 5 per group. Data are presented as mean ± s.e.m. from at least three independent experiments. All p-values were calculated using one-way ANOVA. Statistical significance is indicated as follows: \**p*<0.05, \*\**p*<0.01, \*\*\**p*<0.001 and \*\*\*\**p*<0.0001 vs. indicated group; ns, not significant.

### MLCK-dependent mechanisms underlie small intestinal barrier disruption in colitis

Given that certain inflammatory cytokines are known to increase paracellular permeability by upregulating the expression and activity of the long myosin chain kinase (MLCK) isoform (Al-Sadi et al., 2008; Al Nabhani et al., 2017b; Jung et al., 2012; Wang et al., 2005; Wang et al., 2006), we investigated to which extent the distinct immune responses in TNBS-, DSS-, and oxazolone-induced colitis models influence MLCK (*Mylk*) mRNA expression in the small intestine. Interestingly, the duodenum and ileum of mice from all colitis models exhibited significantly higher *Mylk* mRNA expression compared to non-colitic control mice (*Fig. 2A*). This upregulation suggests that elevated MLCK expression may contribute to the increased intestinal permeability observed across these colitis models, independent of the specific immune response profile. To further explore the role of MLCK, we assessed whether its pharmacological inhibition with ML7 (an inhibitor of MLCK) affects the type of immune responses in the small intestine under colitis induction. While ML7 treatment had minimal impact on body weight in TNBS-, DSS-, and oxazolone-treated mice (data not shown), it led to a reduction in disease activity index (DAI), colon shortening, and macroscopic damage scores (*Fig. S2 A–C*). Moreover, MLCK inhibition significantly suppressed the overexpression of proinflammatory cytokines TNF-α, IFN-γ, IL-1β, and IL-12 in the duodenum and ileum across all three colitis models (*Fig. 2B* and *Fig. S2 D*). In contrast, ML7 had no effect on the elevated expression of Th2-associated cytokines, IL-4, IL-5, and IL-13, observed in the small intestine of oxazolone-treated mice (*Fig. 2C* and *Fig. S2 E*). These findings highlight a divergence in the regulation of Th1-versus Th2-mediated inflammatory responses in colitis and underscore MLCK’s selective involvement in modulating Th1-driven small intestinal inflammation and barrier dysfunction. In addition, MLCK inhibition did not alter *Tnfr1* mRNA expression but significantly reduced the colitis-induced upregulation of *Tnfr2*, *Tjp1*, and *Tjp2* mRNA levels in the small intestine, genes associated with barrier dysfunction (*Fig. 2D* and *Fig. S2 D*). Notably, *in vivo* and *ex vivo* assessments of paracellular permeability in ML7-treated mice showed values comparable to those observed in non-colitic controls (*Fig. 2E–F*), indicating effective restoration of barrier integrity. Furthermore, treatment with ML7 led to a downregulation of *Mylk* mRNA expression across all three colitis models (*Fig. 2G*), further supporting the role of MLCK in mediating small intestinal permeability defects during colonic inflammation. To further confirm the involvement of MLCK in intestinal barrier dysfunction, we conducted experiments using mice deficient in the long MLCK isoform (MLCK^KO^). Similar to the effects observed with ML7 treatment, MLCK^KO^ mice developed slightly less severe TNBS-induced colitis (*Fig. S2 F–H*) and were protected from the duodenal and ileal paracellular permeability increases seen in control littermate mice (*Fig. 2H*). Together, these findings show that MLCK plays a critical role in compromising small intestinal integrity during colitis, underscoring its function in mediating permeability defects distant from the primary sites of colonic inflammation.

**Figure 2:**
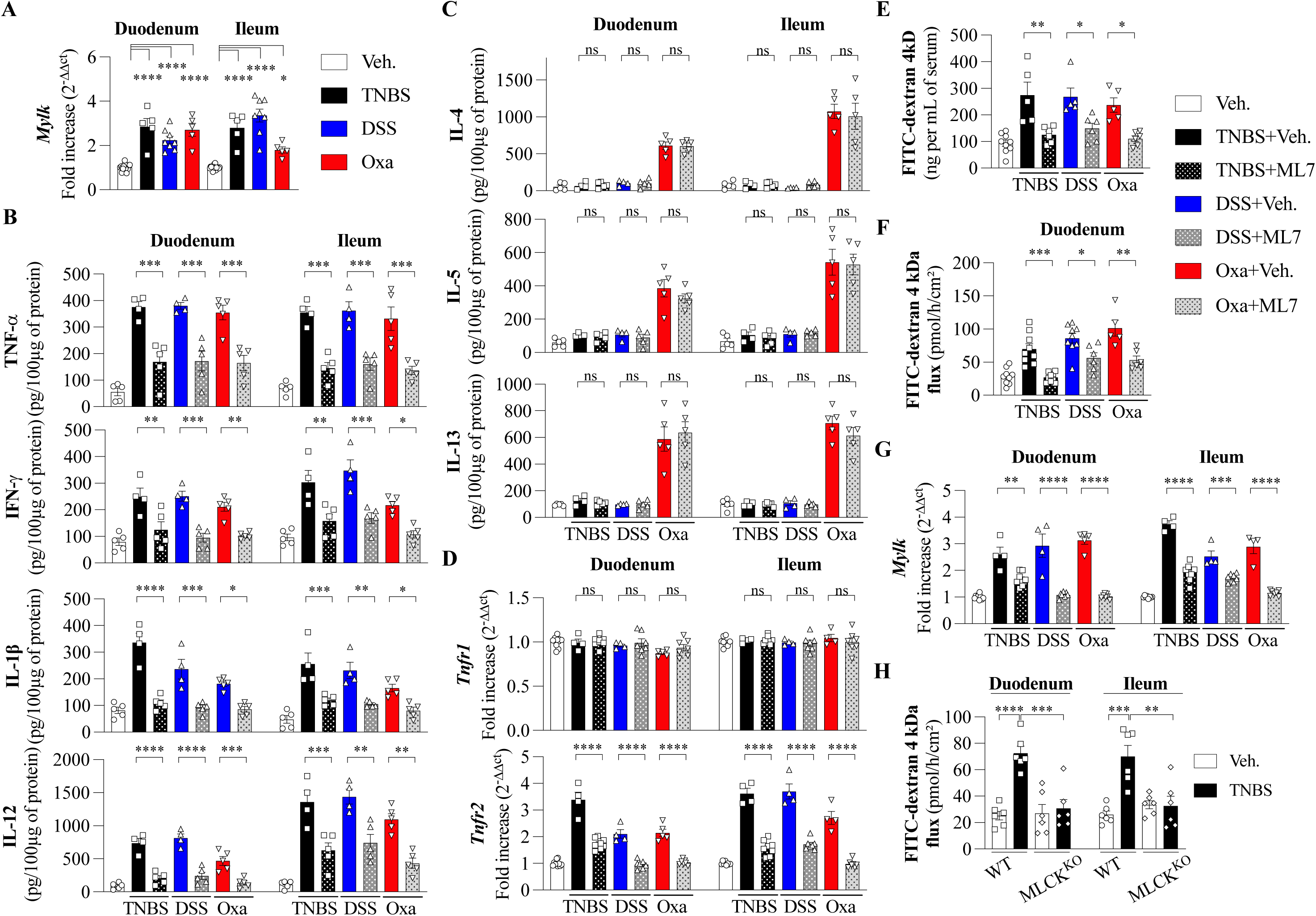
Inhibition of MLCK prevents the increased paracellular permeability of small bowel triggered by colitis. **(A)** mRNA expression of *Mylk* in the duodenum and ileum of WT mice with TNBS-(black), DSS-(blue), or oxazolone-induced (red) colitis. Control mice received vehicle only (white). **(B-G)** Effect of ML7 treatment on cytokine production, barrier function, and gene expression in the duodenum and ileum of WT mice with TNBS-, DSS-, or oxazolone-induced colitis. Control mice received vehicle only. **(B and C)** Cytokine production measured by ELISA. **(B)** TNFα, IFNγ, IL-1β, IL-12 and **(C)** IL-4, IL-5, and IL-13 protein levels. **(D)** *Tnfr1* and *Tnfr2* mRNA expression. **(E)** Gut paracellular permeability assessed by oral FITC-dextran gavage. **(F)** Duodenal paracellular permeability measured by Ussing chamber. **(G)** *Mylk* mRNA expression following ML7 treatment versus vehicle controls. **(H)** Duodenal and ileal paracellular permeability in TNBS-induced colitis, comparing WT (white) and MLCK-knockout mice (black) measured by Ussing chamber. Each data point represents one mouse; *n* ≥ 4 per group. Data are presented as mean ± s.e.m. from at least three independent experiments. All p-values were calculated using one-way ANOVA. Statistical significance is indicated as follows: \**p*<0.05, \*\**p*<0.01, \*\*\**p*<0.001 and \*\*\*\**p*<0.0001 vs. indicated group; ns, not significant.

### Colitis-induced immune responses differentially affect small intestinal function via microbiota-dependent mechanisms

Given the pivotal role of the intestinal microbiota in regulating gut homeostasis, we investigated it influence on paracellular permeability and small intestinal function in the context of colitis. Thus, broad-spectrum antibiotics (ABX) were administered via drinking water during TNBS- or oxazolone-induced colitis in mice. ABX reduce the disease activity index (DAI), colon shortening, and macroscopic damage scores in TNBS-, and oxazolone-treated mice compared to control groups (*Fig. S3 A–B*). We observed that the overexpression of *Mylk* in both TNBS- and oxazolone-treated groups was mediated by microbiota-dependent mechanisms (*Fig. 3A*). This was accompanied by microbiota-dependent regulation of barrier-related genes *Tjp1* and *Tjp2* in the small intestine, which was normalized following antibiotic treatment (*Fig. 3B–C*). Consistently, the enhanced paracellular permeability observed in colitic mice was significantly reduced upon microbiota depletion, as assessed *in vivo* (*Fig. 3D*) and *ex vivo* using Ussing chamber (*Fig. 3E*). In parallel, *Tnfr2* mRNA expression, but not *Tnfr1*, previously elevated in response to colitis, was also normalized following antibiotic treatment (*Fig. 3F*). These microbiota-dependent changes in barrier function were associated with a reduction in pro-inflammatory cytokine expression. Specifically, antibiotic treatment led to decreased levels of Th1-associated cytokines (TNF-α, IFN-γ, IL-1β, IL-12) in the small intestine of TNBS-treated mice (*Fig. 3G*), and in oxazolone-treated mice, it also suppressed both Th1 and Th2 cytokines (IL-4, IL-5, IL-13) (*Fig. 3G* and *Fig. S3 C–D*), further linking microbial regulation to immune modulation and barrier integrity. Taken together, these data demonstrated that alteration of small bowel physiology induced by colitis is modulated by gut microbiota.

**Figure 3:**
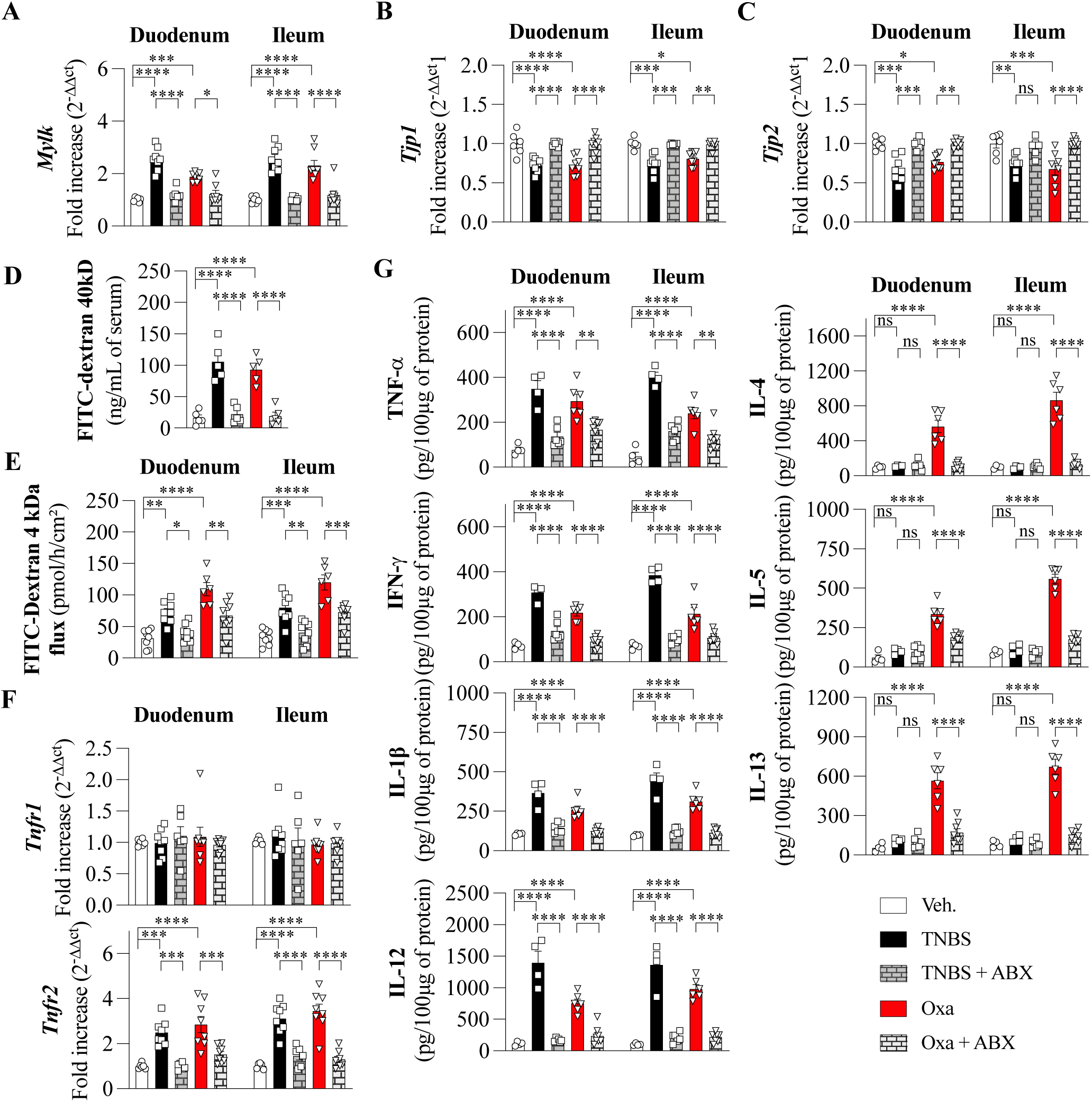
The increased paracellular permeability of small bowel triggered by colitis is dependent of microbiota. **(A-G)** Effect of antibiotic treatment on cytokine production, barrier function, and gene expression in the duodenum and ileum of WT mice with TNBS-, or oxazolone-induced colitis. Control mice received vehicle only. **(A-C)** mRNA expression of **(A)** *Mylk*, **(B)** *Tjp1* and **(C)** *Tjp2* in the duodenum and ileum of WT mice. **(D)** Gut paracellular permeability assessed by oral FITC-dextran gavage. **(E)** Duodenal paracellular permeability measured by Ussing chamber. **(F)** *Tnfr1* and *Tnfr2* mRNA expression **(G)** Cytokine production of TNFα, IFNγ, IL-1β, IL-12, IL-4, IL-5, and IL-13 protein levels measured by ELISA. One data point represents one mouse; at least *n* = 4 per group. Data are presented as mean ± s.e.m. from at least three independent experiments. All *p-*values were calculated using one-way ANOVA. Statistical significance is indicated as follows: \**p*<0.05, \*\**p*<0.01, \*\*\**p*<0.001 and \*\*\*\**p*<0.0001 vs. indicated group; ns, not significant.

### RORγt⁺ CD4⁺ T cells regulate small intestinal permeability in TNBS-but not oxazolone-induced colitis

To investigate the role of CD4⁺ T cell subsets in colitis-induced changes in small intestinal homeostasis, we examined immune cell composition in the small intestinal *lamina propria* of mice with colitis induced by TNBS, DSS, or oxazolone. Flow cytometry analysis revealed a significant increase in RORγt⁺ Foxp3⁻ CD4⁺ T cells (Th17 cells) in the small intestine of TNBS- and DSS-induced colitic mice compared to vehicle controls (*Fig. 4A–B*). This expansion was characteristic of the Th1/17-skewed immune response seen in these colitis models. However, no significant increase in Gata3⁺ Foxp3⁻ CD4⁺ T cells (Th2 cells) was observed in any of the models (*Fig. 4A–B*). Interestingly, Th17 cell expansion was absent in the oxazolone-induced colitis model. Additionally, we did not observe any significant alterations in the regulatory Foxp3⁺ CD4⁺ T cell (Treg) populations in the small intestine, suggesting that Th17 cells, rather than Tregs or Th2 cells, play a major role in colitis-induced immune responses in the small intestine in TNBS and DSS colitis model. Importantly, the expansion of Th17 cells in the small intestine was microbiota-dependent since antibiotic treatment significantly reduced the frequency of RORγt⁺ Foxp3⁻ CD4⁺ T cells in colitic mice, indicating that the presence of the microbiota is essential for the observed increase in Th17 cells in the small intestine (*Fig. 4C*). These findings suggest that Th17 cell expansion in response to TNBS and DSS colitis is driven by the microbiota, reinforcing the pivotal role of microbiota in regulating immune responses in the gut.

**Figure 4:**
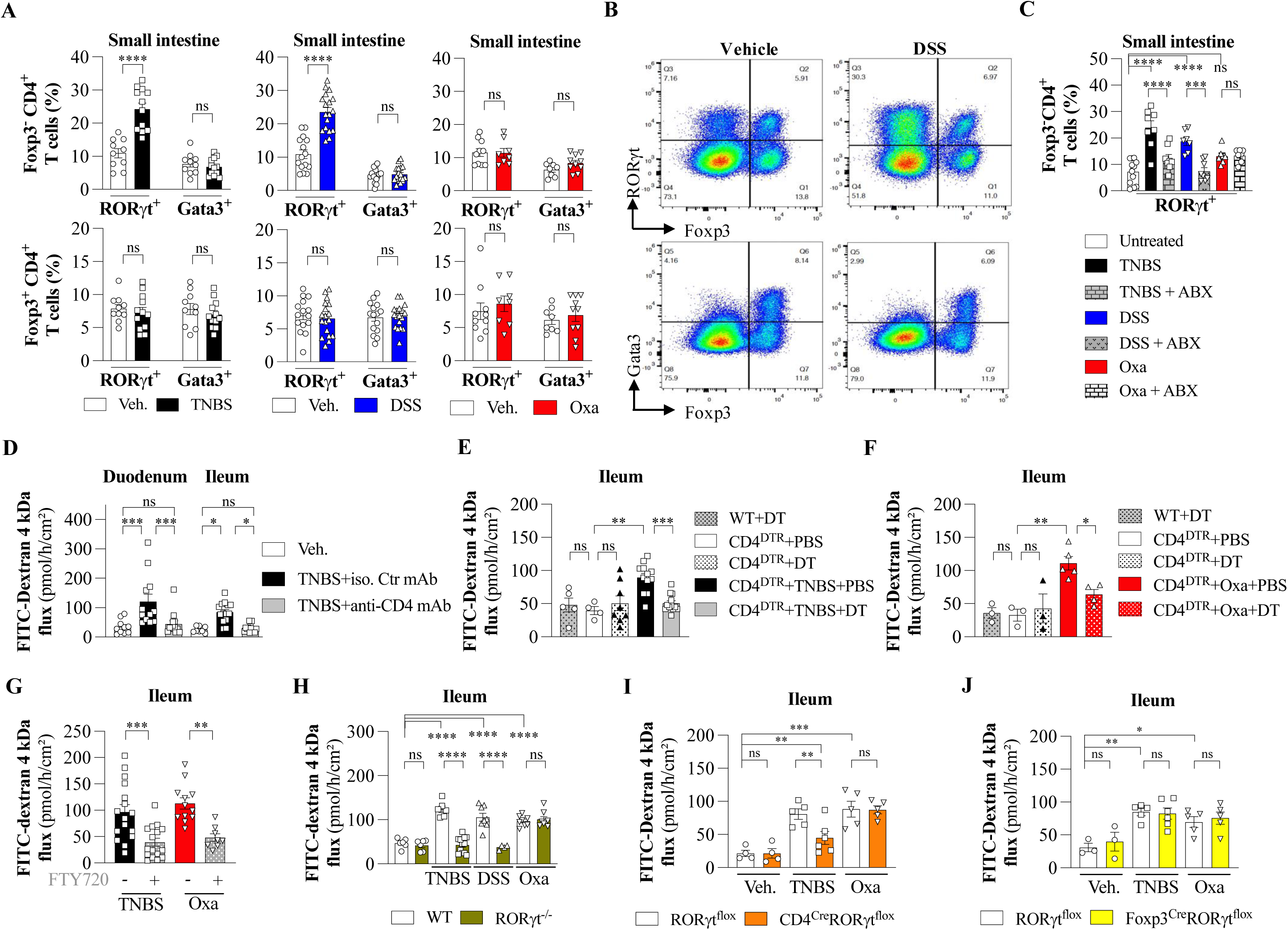
Increased paracellular permeability of small intestine is mediated by re-circulating CD4^+^ T-cells from inflamed colon to small intestine. **(A)** Percentages of CD4⁺ T cell subsets expressing RORγt and Gata3 in Foxp3⁻ or Foxp3^+^ CD4⁺ T cells isolated from the small intestine following colitis induction with TNBS, DSS, or oxazolone. **(B)** Representative flow cytometry plots of CD4⁺ T cells expressing RORγt, Foxp3, and Gata3 isolated from the small intestine of DSS-treated colitic mice and vehicle-treated controls. **(C)** Frequency of RORγt⁺Foxp3⁻CD4⁺ T cells in the small intestine of control and antibiotics (ABX)-treated WT mice following induction of colitis with TNBS, DSS, or oxazolone. **(D)** Duodenal and ileal paracellular permeability assessed by Ussing chamber in WT mice following CD4⁺ T cell depletion by monoclonal antibody or isotype control after induction of TNBS colitis. **(E-J)** Ileal paracellular permeability in: CD4^DTR^ mice treated with diphtheria toxin (DT) to deplete CD4⁺ T cells following TNBS **(E)** or oxazolone **(F)** colitis induction versus WT control mice treated with DT or PBS vehicle; **(G)** WT mice treated with FTY720 (a CD4⁺ T cell recirculation inhibitor) or vehicle; **(H)** RORγt^−/-^ mice and WT littermate controls following TNBS or oxazolone-induced colitis; **(I)** CD4^Cre^RORγt^flox^ mice and RORγt^flox^ littermate controls; and **(J)** Foxp3^Cre^RORγt^flox^ mice and RORγt^flox^ littermate controls, all following TNBS or oxazolone-induced colitis. Each data point represents one mouse; *n* ≥ 3 per group. Data are presented as mean ± s.e.m. from at least three independent experiments. All *p-*values were calculated using one-way ANOVA. Statistical significance is indicated as follows: \**p* < 0.05, \*\**p* < 0.01, \*\*\**p* < 0.001, \*\*\*\**p* < 0.0001 vs. indicated group; ns, not significant.

The increased paracellular permeability observed in the small intestine of colitic mice was primarily driven by CD4⁺ T cells. Depletion of CD4⁺ T cells using monoclonal antibodies significantly reduced DAI, Wallace scores, and colonic shortening (*Fig. S4 A and D*) and duodenal and ileal permeability, as assessed by Ussing chamber analysis, in TNBS-induced colitic mice (*Fig. 4D*). Additionally, in the CD4^DTR^ mouse model, where CD4⁺ T cells are selectively depleted upon administration of diphtheria toxin (DT), DAI, Wallace scores, and colonic shortening (*Fig. S4 B,C and E,F*) and ileal paracellular permeability were markedly reduced after colitis induction with TNBS (*Fig. 4E*) and oxazolone (*Fig. 4F*). These findings indicate that the increased intestinal permeability in the small intestine during colitis is largely mediated by CD4⁺ T cells, supporting the hypothesis that immune cell subsets, particularly Th17 cells, play a critical role in disrupting small intestinal barrier function. To further investigate the role of CD4⁺ T cell recirculation in the ileal increased paracellular permeability during colitis, we utilized FTY720, a sphingosine-1-phosphate receptor modulator that inhibits lymphocyte egress from lymphoid organs. FTY720 treatment partially ameliorated colitis symptoms (data not shown) and reduced ileal permeability in both TNBS- and oxazolone-induced colitis models (*Fig. 4G*). This suggests that CD4⁺ T cell recirculation, and not merely their presence in the small intestine, is crucial for mediating the intestinal barrier dysfunction observed in these models.

As it has been reported that the absence of RORγt in mice results in increased susceptibility to colitis, characterized by a more severe inflammatory response and greater tissue damage, we found similar outcomes in our study. DAI, Wallace scores, and colonic shortening were significantly higher in RORγt-deficient mice compared to WT mice across all three colitis models (*Fig. S4 G–K*). While colonic permeability was similarly elevated in both genotypes (*Fig. S4 L*), ileal permeability was differentially affected where RORγt-deficient mice exhibited reduced paracellular permeability in the ileum following TNBS or DSS treatment but not oxazolone (*Fig. 4H*). This distinction suggests that small intestinal barrier function is regulated by RORγt⁺ T cells in a context-dependent manner, influenced by the nature of the immune response induced by each colitis model. To confirm the specific contribution of RORγt⁺ CD4⁺ T cells to small intestinal barrier dysfunction, we used CD4^Cre^RORγt^flox^ mice, in which RORγt is selectively deleted in CD4⁺ T cells (*Fig. 4I*). In this model, we observed a significant reduction in DAI, Wallace scores, and colonic shortening (*Fig. S5*) and ileal paracellular permeability following TNBS-induced colitis, comparable to that seen in global RORγt-deficient mice. In contrast, DAI, Wallace scores, and colonic shortening (*Fig. S5*) and ileal permeability remained elevated in CD4^Cre^RORγt^flox^ mice subjected to oxazolone-induced colitis, similar to control littermates (*Fig. 4I*). These results indicate that RORγt expression in CD4⁺ T cells contributes to increased small intestinal permeability specifically in the context of TNBS-induced inflammation, and is dispensable under Th2-polarized conditions such as those elicited by oxazolone. When RORγt was selectively deleted in Foxp3⁺ Tregs using the Foxp3^Cre^RORγt^flox^ model, ileal paracellular permeability remained elevated following both TNBS- and oxazolone-induced colitis, comparable to control littermates (*Fig. 4J* and *Fig. S5*). This suggests that RORγt expression in Tregs is not required for the regulation of small intestinal barrier function in the context of colonic inflammation. Instead, RORγt⁺ conventional CD4⁺ T cells, rather than Tregs, appear to be the primary drivers of the increased permeability observed in the small intestine during Th1/Th17-skewed colitis.

Altogether, these findings suggest that the contribution of RORγt⁺ CD4⁺ T cells to paracellular permeability and small bowel barrier dysfunction is context-dependent and varies with the type of immune response elicited by different colitis models.

### Upregulation of Nod2 in the small intestine is model-specific and modulated by CD4⁺ T Cells, MLCK activity, and microbiota

The induction of colitis by TNBS or DSS in adult WT mice led to a significant increase in *Nod2* mRNA expression in the small intestine, even in regions distant from the primary site of inflammation. This upregulation was not observed in oxazolone-treated mice, indicating that *Nod2* induction is specific to TNBS and DSS colitis models (*Fig. 5A*).

**Figure 5:**
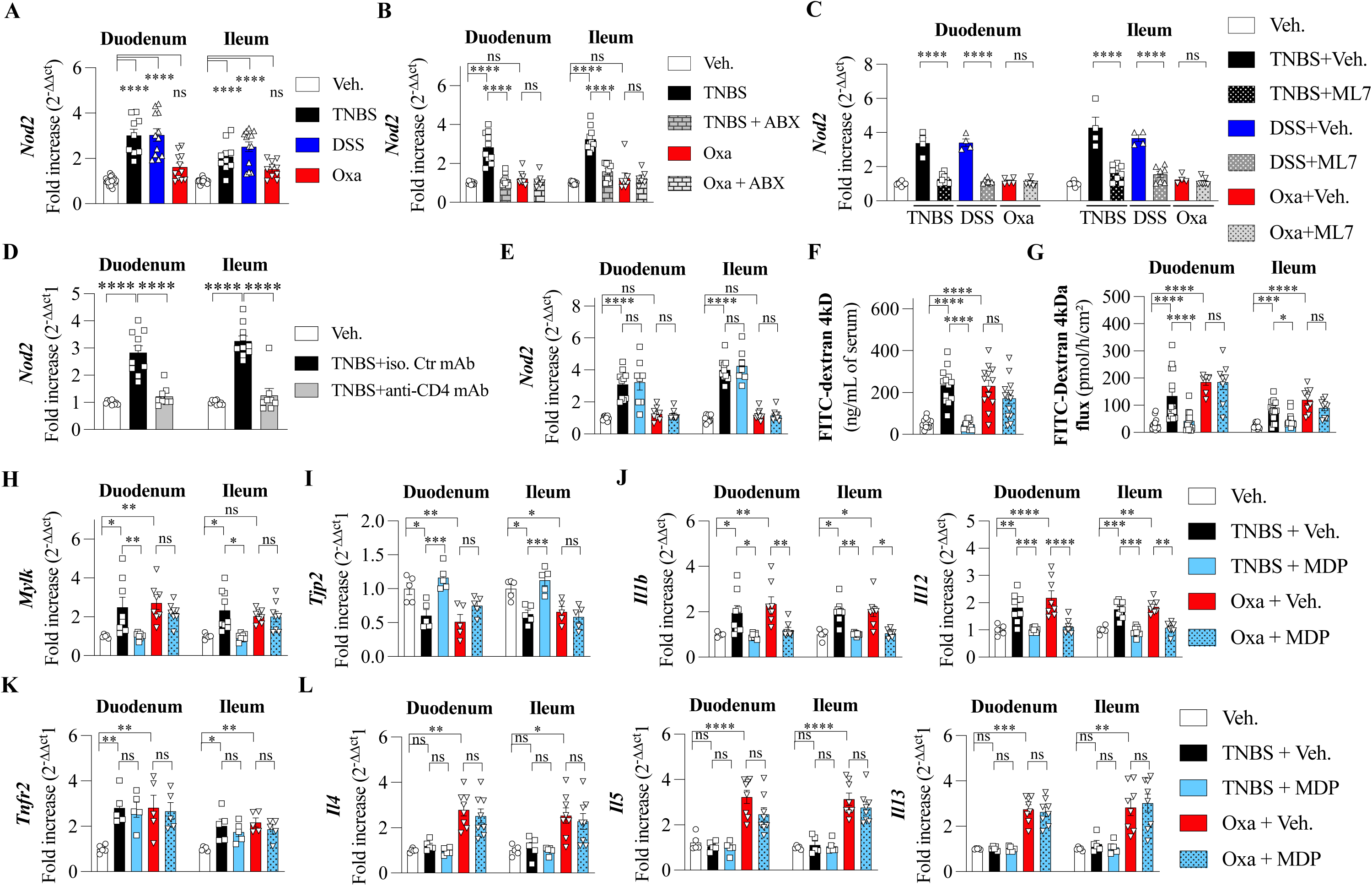
Nod2 stimulation suppresses the increased paracellular permeability of the small intestine induced by colitis. **(A)** duodenal and ileal *Nod2* mRNA expression in WT mice with colitis induced by TNBS (black), DSS (blue), or oxazolone (red) or control mice received vehicle only (white). **(B)** *Nod2* expression in TNBS- or oxazolone-induced colitic mice treated or not with antibiotics (ABX). **(C)** *Nod2* expression in TNBS-, DSS-, or oxazolone-induced colitic mice treated or not with ML7, an inhibitor of MLCK. **(D)** *Nod2* expression in TNBS-induced colitic mice following CD4⁺ T cell depletion by monoclonal antibody or isotype control. **(E-L)** Effect of muramyl dipeptide (MDP) treatment on cytokine production, barrier function, and gene expression in the duodenum and ileum of WT mice with TNBS- or oxazolone-induced colitis. Control mice received vehicle only. **(E)** *Nod2* mRNA expression measured by qPCR. **(F)** Paracellular permeability measured by oral FITC-dextran gavage. **(G)** Duodenal and ileal paracellular permeability measured by Ussing chamber, and **(F)** *Mylk*, **(I)** *Tjp2*, **(J)** *Il1b*, *Il12* and **(K)** *Tnfr2* and **(L)** *Il4*, *Il5*, and *Il13* mRNA expression measured by qPCR. Each data point represents one mouse; *n* ≥ 4 per group. Data are presented as mean ± s.e.m. from at least three independent experiments. All *p-*values were calculated using one-way ANOVA. Statistical significance is indicated as follows: \**p*<0.05, \*\**p*<0.01, \*\*\**p*<0.001 and \*\*\*\**p*<0.0001 vs. indicated group; ns, not significant.

Through a series of experiments, we demonstrated that *Nod2* overexpression in the small intestine of TNBS-treated mice is mediated by microbiota-dependent (*Fig. 5B*) and MLCK-dependent mechanisms (*Fig. 5C*). Notably, CD4⁺ T cell depletion using monoclonal antibodies restored *Nod2* expression in the duodenum and ileum of TNBS-treated mice to levels comparable to untreated controls (*Fig. 5D*), suggesting a critical role for CD4⁺ T cells in regulating Nod2 expression during colitis.

Stimulation of Nod2 with muramyl dipeptide (MDP), its canonical ligand, did not affect *Nod2* mRNA levels in the duodenum or ileum of mice with TNBS- or oxazolone-induced colitis (*Fig. 5E*). However, MDP treatment significantly reduced paracellular permeability in the small intestine of TNBS-treated mice, but not in oxazolone-treated mice, as measured *in vivo* (*Fig. 5F*) and *ex vivo* using Ussing chambers (*Fig. 5G*). This improvement in barrier function was associated with a reduction in *Mylk* and *Tjp2* mRNA expression in the small intestine of TNBS-treated mice (*Fig. 5H–I*), suggesting that MDP signaling modulates epithelial integrity through tight junction regulation in a model-dependent manner. Furthermore, MDP administration reduced *Il1b* and *Il12* mRNA expression in the small intestine in both TNBS and oxazolone models (*Fig. 5J*), but did not alter *Tnfr2* expression (*Fig. 5K*). Finally, in oxazolone-induced colitis, the elevated expression of Il4, Il5, and Il13 in the duodenum and ileum was unaffected by MDP treatment, reinforcing the Th2-dominant immune profile of this model and its independent to Nod2-mediated modulation (*Fig. 5L*).

### Hematopoietic and non-hematopoietic roles of Nod2 in small intestinal barrier integrity and inflammation in TNBS-induced Colitis

In patients with CD, *NOD2* is often mutated with the 3020insC mutation encoding a truncated protein (1007fs) being the most deleterious. We therefore chose to study the consequences of a mutated Nod2 protein, 2939insC, the murine equivalent of the mutation found in humans. In both Nod2 knockout (Nod2^KO^) and Nod2^2939insC^ mice with TNBS-induced colitis, small intestinal paracellular permeability was elevated compared to wild-type (Nod2^WT^) mice. While co-treatment with MDP significantly reduced small intestinal permeability in Nod2^WT^ mice, no such effect was observed in Nod2^KO^ or Nod2^2939insC^ mice, indicating that the protective effect of MDP requires functional Nod2 (*Fig. 6A*).

**Figure 6:**
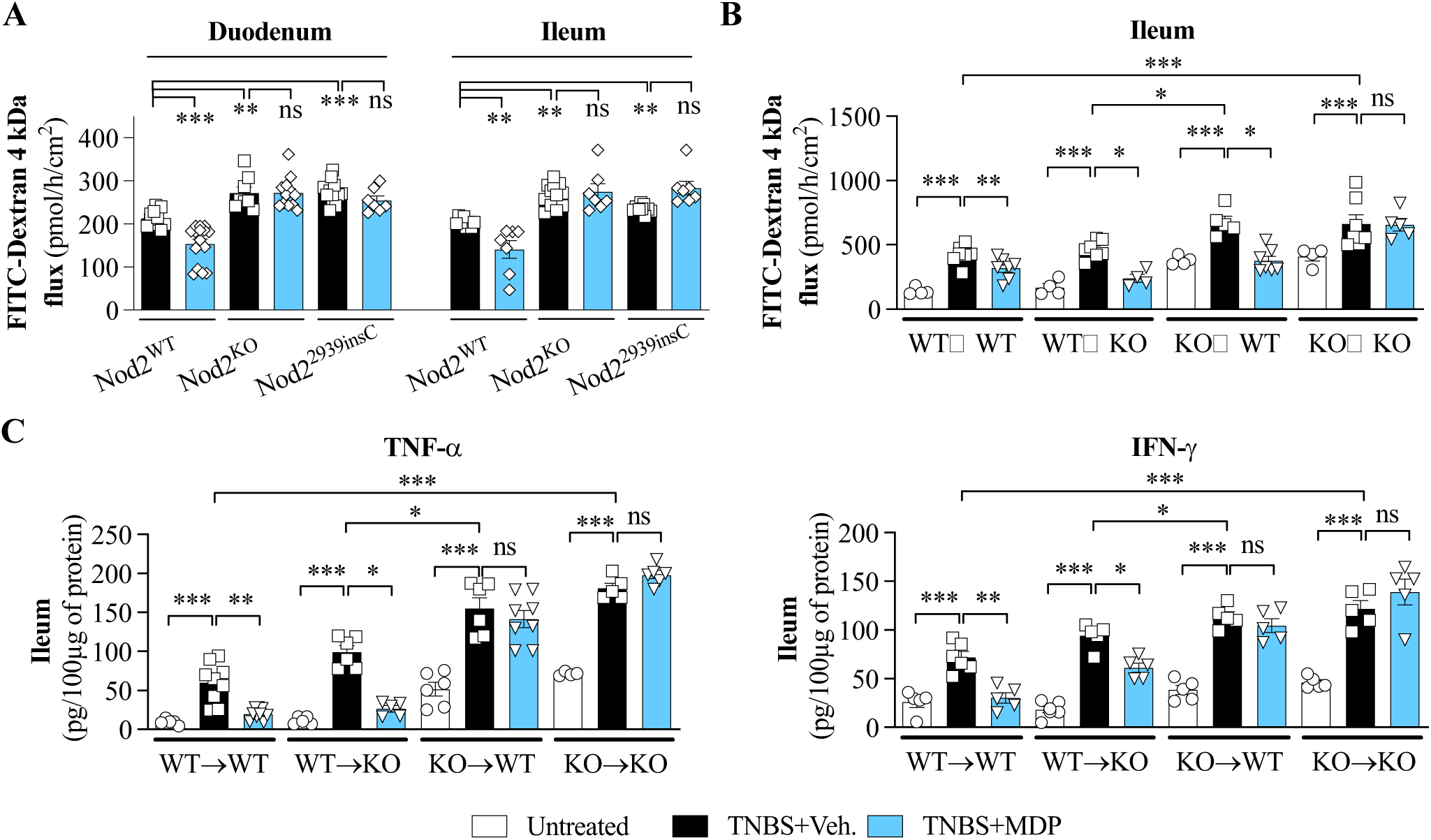
Activation of Nod2 in hematopoietic but not non-hematopoietic *Nod2* reduces the severity of colitis induced by TNBS. **(A)** Duodenal and ileal paracellular permeability measured by Ussing chamber in *Nod2*^WT^*, Nod2*^KO^ and *Nod2*^2939insC^ mice following TNBS-induced colitis. Mice were treated with muramyl dipeptide (MDP) or PBS (vehicle), as indicated. **(B-C)** Chimeric mice were generated by transplantation of bone marrow stem cells (BMSCs) from *Nod2*^WT^ to *Nod2*^KO^ (WT→KO) or from *Nod2*^KO^ to *Nod2*^WT^ (KO→WT). Mice receiving BMSCs from donors of the same genotype (WT→WT or KO→KO) served as controls. **(B)** Duodenal and ileal paracellular permeability measured by Ussing chamber. **(C)** Cytokine levels in inflamed colonic tissue. Each data point represents one mouse; *n* ≥ 4 per group. Data are presented as mean ± s.e.m. from at least three independent experiments. All *p-* values were calculated using one-way ANOVA. Statistical significance is indicated as follows: \**p*<0.05, \*\**p*<0.01, \*\*\**p*<0.001 and \*\*\*\**p*<0.0001 vs. indicated group; ns, not significant.

Since Nod2 is expressed in both hematopoietic and non-hematopoietic compartments, we employed a bone marrow chimera model to dissect the compartment-specific role of Nod2 in regulating small intestinal barrier function during TNBS-induced colitis. Bone marrow stem cells (BMSCs) were transplanted from Nod2^KO^ mice into Nod2^WT^ recipients (KO→WT) and vice versa (WT→KO). These experimental groups were compared to control chimeras receiving BMSCs from donors of the same genotype (WT→WT and KO→KO). Three months post-transplantation, mice were challenged with TNBS. Following TNBS induction, paracellular permeability (*Fig. 6B*), IFN-γ and TNF-α (*Fig. 6C*) in ileum were significantly increased in mice lacking Nod2 in the hematopoietic compartment (KO→WT and KO→KO), highlighting the contribution of hematopoietic *Nod2* to the regulation of intestinal inflammation and barrier integrity. Importantly, MDP treatment ameliorated small intestinal inflammation and reduced permeability in chimeric mice that expressed Nod2 in the hematopoietic compartment, compared to mice reconstituted with Nod2^KO^ bone marrow (Fig. 6B–C and Fig. S6). Conversely, Nod2 expression in the non-hematopoietic compartment was sufficient to reduce excess small intestinal permeability, but not pro-inflammatory cytokine expression, following MDP treatment, regardless of the Nod2 status in hematopoietic cells. Together, these findings indicate that Nod2 confers protective effects on the gut barrier via both hematopoietic and non-hematopoietic compartments. While non-hematopoietic Nod2 appears sufficient to preserve barrier integrity, hematopoietic Nod2 is necessary to limit pro-inflammatory cytokine responses in the small intestine during colitis.

In addition, we investigated the link between excessive intestinal permeability and the development of inflammatory lesions in *Nod2* deficient (*Nod2*^KO^) mice (*Fig. 7*). To this end, *Nod2*^KO^ mice were treated with ML7, and both small intestinal permeability and the progression of inflammatory lesions were assessed. ML7 treatment significantly attenuated the development of inflammatory lesion formation in the small intestine (*Fig. 7A–B*), prevented villus shortening (Figure 7C), and preserved goblet cells (*Fig. 7D*). Moreover, ML7 treatment reduced immune cell infiltration along the villus axis (*Fig. 7E–F*) and mitigated the excessive paracellular permeability typically observed in *Nod2*^KO^ mice (*Fig. 7G*).

**Figure 7:**
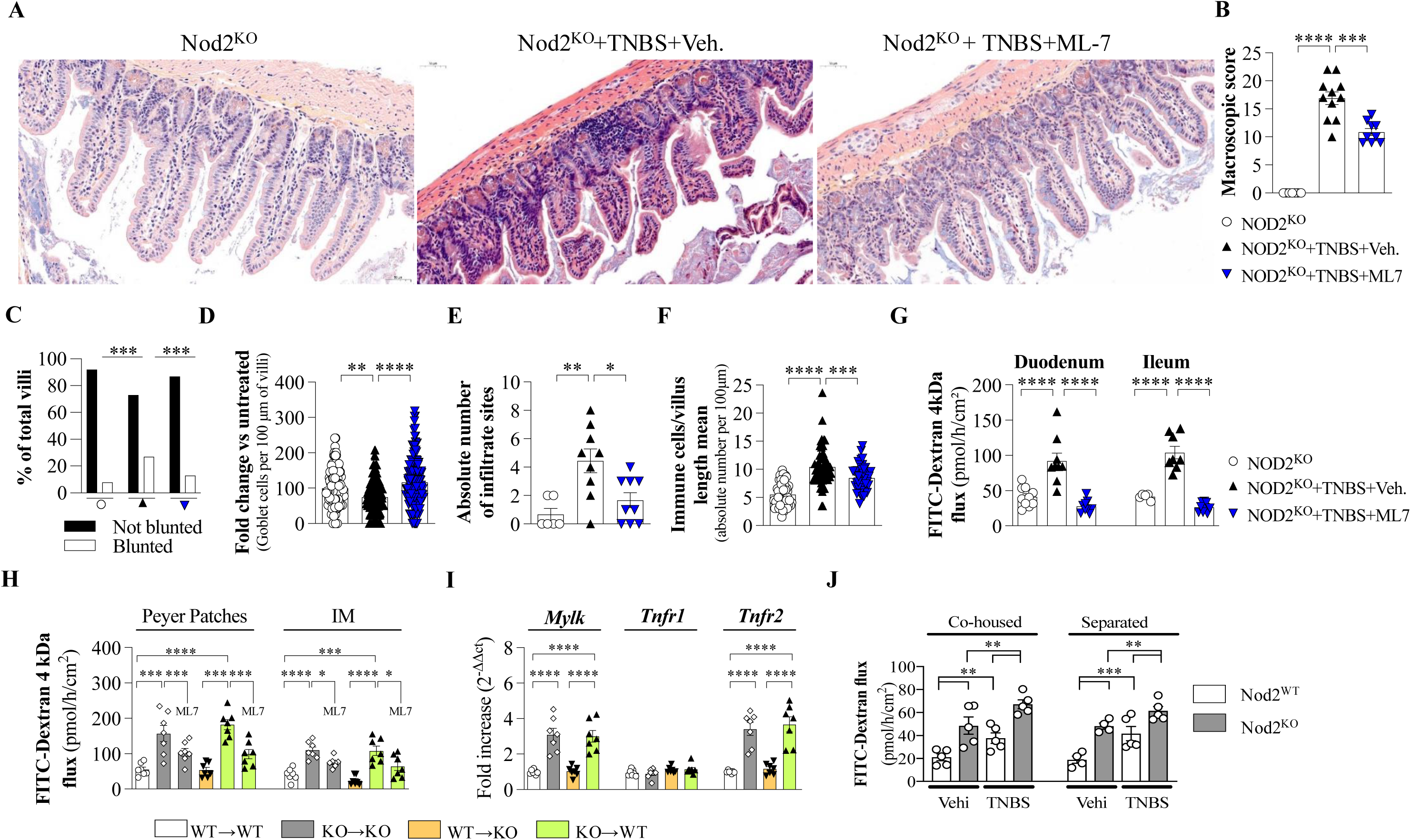
The gut dysbiosis linked to Nod2 deletion does not contribute to the increased permeability. **(A-G)** Nod2-deficient (*Nod2*^KO^) mice following TNBS-induced colitis, treated with either ML-7 or vehicle control, compared to untreated control mice. **(A)** Representative histological sections of the small intestine. **(B)** Microscopic inflammation scores. **(C)** Percentage of villi exhibiting blunting versus normal morphology. **(D)** Goblet cell counts per 100 µm of villus length. **(E)** Absolute number of inflammatory infiltrate sites per section. **(F)** Number of immune cells normalized to villus length. **(G)** Duodenal and ileal paracellular permeability measured by Ussing chamber. **(H-I)** Chimeric mice were generated by transplantation of bone marrow stem cells (BMSCs) from *Nod2*^WT^ to *Nod2*^KO^ (WT→KO) or from *Nod2*^KO^ to *Nod2*^WT^ (KO→WT). Mice receiving BMSCs from donors of the same genotype (WT→WT or KO→KO) served as controls. **(H)** Peyer Patches and ileal paracellular permeability measured by Ussing chamber. **(I)** Ileal *Mylk, Tnfr1 and Tnfr2* mRNA expression measured by qPCR. **(J)** Ileal paracellular permeability measured by Ussing chamber in *Nod2*^KO^ and *Nod2*^KO^ mice that were either cohoused (maintained in the same cage) or separated (housed in different cages) after weaning, followed by TNBS-induced colitis in adulthood. Each data point represents one mouse; *n* ≥ 4 per group. Data are presented as mean ± s.e.m. from at least three independent experiments. All *p-*values were calculated using one-way ANOVA. Statistical significance is indicated as follows: \**p*<0.05, \*\**p*<0.01, \*\*\**p*<0.001 and \*\*\*\**p*<0.0001 vs. indicated group; ns, not significant.

To determine whether the protective effect of ML7 was mediated through the hematopoietic or non-hematopoietic compartment, we generated bone marrow chimeric mice with Nod2 deficiency restricted to either the immune or non-immune compartments. We found that ML7 treatment was effective only in chimeric mice lacking Nod2 in the hematopoietic compartment, but not in those lacking Nod2 exclusively in non-hematopoietic cells (*Fig. 7H– I*). In these mice, increased paracellular permeability was reduced, along with the downregulation of mRNA expression levels of *Mylk* and Tnfr2, but not Tnfr1, in the small intestine (*Fig. 7H–I*). These results suggest that the impact of MLCK inhibition in Nod2 deficient mice is mediated through the hematopoietic compartment, emphasizing a critical role for CD4^+^ T cells in regulating small intestinal barrier function and inflammation.

*Nod2*^KO^ mice are known to present a gut microbiota dysbiosis involved in both inflammatory and carcinogenic processes. Thus, we wanted to understand the impact of dysbiosis in the elevated permeability found in the small bowel in the context of colitis. As reported in literature, cohousing allows the transmission of microbiota dysbiosis from *Nod2*^KO^ mice to *Nod2*^WT^ mice. For this purpose, following weaning, young WT and *Nod2*^KO^ mice were cohoused for 6 weeks (Al Nabhani et al., 2016). Transferring the dysbiotic microbiota found in *Nod2*^KO^ mice to WT mice had no impact on the TNBS-induced colitis, either in terms of intestinal permeability (*Fig. 7J*), colonic length or Wallace score (*Fig. S7A-B*). These results proved that the microbiota does not play a role in the alteration of the permeability of the small bowel in the context of colitis.

## Discussion

The IBD, including CD and UC, are characterized by impaired intestinal barrier function, which can extend beyond the sites of primary inflammation. Here, we have demonstrated that colonic inflammation, regardless of its immune polarization, induces a systemic disruption of the small intestinal barrier, as seen in three widely used murine models of colitis: TNBS, DSS, and oxazolone. This work builds on prior findings by illustrating that colonic inflammation does not remain confined to the colon but has systemic effects, especially on the small intestine. To the best of our knowledge, the impact of colitis on the paracellular permeability of the small intestine had never been demonstrated in mice model of colitis (Blair et al., 2006; Du et al., 2016; Hering et al., 2012; Jin and Blikslager, 2020; Long et al., 2018; Mankertz et al., 2009; Nemoto et al., 2011; Shen et al., 2006; Tomita et al., 2008).

Our findings corroborate earlier studies showing that colitis-induced inflammation can affect distant segments of the gastrointestinal tract (Amit-Romach et al., 2006; Barada et al., 2006) observed similar alterations in intestinal permeability due to colonic injury, specifically showing changes in the intestinal epithelium even in the absence of overt histological lesions. Here, we show that TNBS, DSS, and oxazolone colitis all led to significant increases in small intestinal permeability, which were associated with MLCK upregulation and the alteration of tight junction proteins, namely Mylk, Tjp1, and Tjp2. These observations align with Blair et al. (Blair et al., 2006), who suggested that MLCK plays a pivotal role in the regulation of intestinal barrier permeability under inflammatory conditions.

While previous studies have primarily focused on the colon or localized effects of colitis on the intestinal epithelium, our study presents a novel finding: despite the immune polarization differences between the TNBS (Th1/Th17), DSS (Th1/Th17), and oxazolone (Th2) models, all three models resulted in similar increases in small intestinal permeability. Specifically, we observed that MLCK was upregulated in epithelial cells across all three models, suggesting a converging mechanistic pathway involving MLCK activation. This extends previous work showing that MLCK is a critical regulator of barrier function in various IBD models (Su et al., 2009). More importantly, we show that MLCK inhibition (via ML7) ameliorated small intestinal permeability and reduced expression of tight junction proteins, supporting the idea that MLCK-driven tight junction dysregulation contributes significantly to barrier dysfunction in IBD. This finding is consistent with studies by Long *et al*. (Long et al., 2018), who suggested that MLCK inhibition could selectively block Th1-mediated inflammation-induced barrier damage, but had minimal effect on Th2 cytokines, suggesting that MLCK contributes more directly to Th1-mediated barrier dysfunction.

Although TNBS is administered in the rectum, it also alters the small intestine without any overt histological lesions in rats suggesting a remote effect of the colitis on the upper intestine (Al Nabhani et al., 2020; Amit-Romach et al., 2006; Barada et al., 2006). In agreement with these works, we did not observed any overt inflammation in the small intestine. However, in accordance with the altered epithelial cells physiology like reduced villous surface, Sucrase Isomaltase and Aminopeptidase activities, density of goblet cells and amounts of mucin 2, we have evidenced that the paracellular permeability of the small intestine was higher in TNBS infused mice than in control mice. These findings may provide a partial explanation or a molecular substrate for the associated small bowel dysfunctions in context of colitis.

Moreover, our results revealed that the inflammatory environment plays a central role in this process (Graham et al., 2019; Wang et al., 2005). We observed increased expression of TNF-α, IFN-γ, IL-1β, IL-12 cytokines in the small intestine in all colitis models, albeit to varying degrees, and inhibition of MLCK led to selective suppression of Th1 cytokines. Th2-associated cytokines (IL-4, IL-5, IL-13) were selectively induced in oxazolone-treated mice, reinforcing the immune specificity of this model. These results highlight the central role of Th1-associated inflammation in driving barrier dysfunction in TNBS and DSS colitis (Graham et al., 2019; Wang et al., 2005). This distinction suggests that Th1 inflammation, rather than Th2, is more directly linked to MLCK-mediated barrier disruption, which is in line with previous works (Al-Sadi et al., 2008; Su et al., 2009; Wang et al., 2006), who found that Th1/Th17 responses strongly affect epithelial permeability compared to Th2-driven inflammation. Interestingly, Th2 cytokines (IL-4, IL-5, IL-13), which were highly elevated in the oxazolone model, did not exhibit the same effect on barrier function as Th1 cytokines. This model, known for its Th2-mediated inflammation, did not show the same degree of small intestinal permeability increase, despite elevated levels of Th2 cytokines. This supports the findings of Kulik *et al*. (2015), where Th2 cytokines were less effective in driving epithelial damage compared to Th1/Th17-associated responses.

A critical immune axis identified in our study is the contribution of CD4⁺ T cells to remote small intestinal injury. Depletion of CD4⁺ T cells in TNBS-and oxazolone-induced colitis markedly reduced permeability and inflammation in the small intestine. We identified RORγt⁺ CD4⁺ T cells (Th17) as key mediators of this process in TNBS and DSS colitis, where Th17 cell expansion was microbiota-dependent and correlated with increased permeability. In contrast, Th2-driven oxazolone colitis did not elicit a similar Th17 response, and barrier dysfunction occurred independently of RORγt expression in CD4⁺ T cells. These findings suggest that different T helper subsets exert model-specific effects on epithelial homeostasis, with Th17 cells mediating small bowel barrier disruption under Th1/Th17 conditions but not in Th2-skewed inflammation. These results underscore the importance of T cell trafficking in driving systemic effects of colitis, supporting previous findings by Nemoto *et al*. (Nemoto et al., 2011), who reported that colitogenic T cells migrate from the inflamed colon to distant regions of the intestine and contribute to epithelial dysfunction. This abnormal crosstalk strongly supports how the anti-inflammatory drugs used in CD (anti-inflammatory drugs, immune-suppressors and anti-TNF-α antibodies diminish the severity of the disease and restore the gut barrier function (Brown et al., 1999; Suenaert et al., 2002). Reciprocally, our data support the importance to develop drugs allowing the normalization of the intestinal barrier function to reduce the inflammatory status.

Our study also identified a crucial protective role for Nod2, a pattern recognition receptor, in maintaining small intestinal barrier function. Nod2 expression was significantly upregulated in the small intestine of TNBS and DSS-treated mice, but not in oxazolone-treated animals, suggesting that Nod2 activation may be particularly important in Th1/Th17-driven colitis. These findings align with our previous observation showed that Nod2 plays a protective role in the gut by modulating epithelial integrity and immune responses (Al Nabhani et al., 2020). Moreover, our Nod2-deficient mice exhibited exacerbated barrier dysfunction and were unresponsive to MDP, supporting our previous studies that Nod2 activation can ameliorate barrier defects in colitis (Al Nabhani et al., 2020; Al Nabhani et al., 2017a) and IBD (Li et al., 2016). Finally, we have demonstrated that Nod2 expression at remote of the colonic lesions had a protective role on both paracellular permeability and secretion of inflammatory cytokines. This beneficial activation of Nod2 may also counteract the effect of IFN-γ and TNF-α suggesting that treatment of patients not carrying mutations in *NOD2* with Nod2 agonists could activate the negative feedback loop to maintain the homeostasis of the permeability then reducing the extension of the inflammatory process (Rosenstiel et al., 2003). In agreement with a previous study, our data evidenced that Nod2 deletion in the hematopoietic compartment is associated with a less colitis severity (Penack et al., 2009). Finally, our data are in agreement with previous study evidencing that some elements of the intestinal barrier are regulated by Nod2 in the hematopoietic compartment like the immune status (Alnabhani et al., 2016) (secretion of cytokines) and the development of inflammatory lesions in the small intestine (Al Nabhani et al., 2020), while the paracellular permeability is controlled by Nod2 in both compartments(Al Nabhani et al., 2017b).

Finally, since microbiota dysbiosis is largely known to alter the intestinal homeostasis in context of colitis (Garrett et al., 2010), we have checked if the microbiota dysbiosis linked to Nod2 deletion is involved or not in the increased paracellular permeability of the small intestine in context of colitis. After transfer of microbiota dysbiosis into WT mice by cohousing condition, we have shown that the increased paracellular permeability of WT-cohoused with *Nod2^KO^* mice is not more important than those observed in WT mice no-cohoused with *Nod2^KO^*. These data are in agreement with previous study reporting that the transfer of microbiota dysbiosis associated with *Nod2^KO^* into WT mice did not affect the gut paracellular permeability of WT (Al Nabhani et al., 2016). However, we did not confirm the increased susceptibility to develop colitis and/or cancer linked to the microbiota dysbiosis linked to Nod2 deletion reported in the study of Couturier-Maillard et al (Couturier-Maillard et al., 2013).

In conclusion, our findings demonstrate that distinct immune pathways in colitis converge on shared effector mechanisms, namely, microbiota-, MLCK-, and CD4⁺ T cell, dependent disruption of small intestinal barrier function. We identify Nod2 as a critical protective factor that operates across hematopoietic and epithelial compartments to maintain barrier integrity. These insights broaden our understanding of systemic effects of localized gut inflammation and suggest that therapeutic strategies aimed at modulating MLCK activity, lymphocyte trafficking, and NOD2 signaling may hold promise in restoring epithelial function in IBD.

## Material and Methods

### Animal models

C57BL/6 wild-type (WT), *Nod2* null allele (*Nod2*^KO^), *Nod2*^2939insC^ (homozygotes for a mutation homologous to the Human 3020insC variant) and *Rorγt*^−/-^ mutant mice were bred or housed in pathogen-free animal facilities (Barreau et al., 2007; Clayburgh et al., 2005; Maeda et al., 2005). *MLCK* deficient mice (MLCK*^KO^)* have been obtained from JT. CD4^Cre^ (JAX stock #022071), Rorγt^flox^ (JAX stock #008771) and Foxp3^Cre^ (JAX stock #016959) mice purchased from Jax (Choi et al., 2016; Lee et al., 2001; Rubtsov et al., 2008). Pathogen-free conditions were monitored every six months in accordance with the full set of FELASA high-standard recommendations. All animals were maintained under 12-hour light–dark cycles with unrestricted access to food and water. Housing and experiments adhered to institutional animal care guidelines and were approved by the local ethical committees for animal experimentation, in accordance with Swiss Federal regulations and the Commission for Animal Experimentation of Kanton Bern. The experimental protocols have been approved by the French government ethical committee for animal experimentation (#28278-2020111810056802).

CD4^+^ T-cells were depleted by two intra-peritoneal (i.p) injections of 100μg purified GK1.5 (anti-L3T4 (CD4^+^) monoclonal antibody (Pharmingen, Germany), 96 and 24 hours before experimentation and 24 hours after TNBS administration (Al Nabhani et al., 2020). The effectiveness of CD4^+^ depletion in Peyer’s plates was checked as previously reported (data not shown)(Al Nabhani et al., 2020). To inhibit the recirculation of CD4^+^ T-cells, mice were treated i.p with FTY720 (3mg/kg; Sigma, France) 0, 1, 2 and 3 days after TNBS infusion (Daniel et al., 2007).

MLCK inhibition was achieved by i.p injection of ML-7, 2 mg/kg body weight (Sigma, France) twice daily during 4 days before experiments and during the TNBS procedure (Barreau et al., 2010).

For the construction of chimeric mice, five million bone marrow stem cells (BMSC) were isolated from WT Ly5.1 or *Nod2*^KO^Ly5.2 mice and injected intravenously either into WT Ly5.1 or *Nod2*^KO^Ly.5.2 lethally-irradiated recipients (Alnabhani et al., 2016; Jung et al., 2012). Chimerism was verified at week 12 by flow cytometry using Ly5.1 and Ly5.2 congenic markers (data not shown) (Al Nabhani et al., 2020).

To investigate the effect of Nod2 stimulation, adult mice were pre-treated i.p with muramyl dipeptide (MDP, 100µg/mice/day; Sigma, France) for 2 consecutive days before experimentation and during the TNBS procedure (Al Nabhani et al., 2020).

The putative impact of *Nod2*-related dysbiosis on the studied phenotypes was assessed using WT and *Nod2^KO^* mice cohoused for 6 weeks in the same cage where indicated (Al Nabhani et al., 2016; Robertson et al., 2013).

Mice were treated during the indicated periods of time with a single antibiotic or a cocktail of antibiotics in their drinking water, containing 1 mg/ml ampicillin, 0.5 mg/ml vancomycin and 0.5 mg/ml metronidazole. The antibiotic-containing drinking water was changed twice per week. Once the treatment was terminated, treated and untreated mice were co-housed.

### Models of colitis

#### TNBS-induced colitis

Colitis was induced in 12-week-old mice by a single intra-rectal administration of 2,4,6-trinitrobenzene sulfonic acid (TNBS, Sigma, France), which was dissolved in ethanol (50:50 vol/vol) at a dose of 120 mg/kg body weight under anaesthesia (Al Nabhani et al., 2020; Barreau et al., 2007). Groups used as controls (vehicle) received an equal volume of PBS and Ethanol (50:50 vol/vol) intra-rectally. A 100 µl aliquot of the freshly prepared solution was injected into the colon, 4 cm from the anus, using a 3.5 F polyethylene catheter as previously described (Barreau et al., 2007). Body weight loss and disease activity index (DAI) were monitored before and 72h after TNBS administration. Mice were sacrificed by cervical dislocation. Colonic length and macroscopic damage Wallace score were recorded (Wallace and Keenan, 1990).

#### DSS-induced colitis

12-week-old mice were exposed to drinking water supplemented with 2.5% Dextran Sulfate Sodium (DSS) for 5 days. Then, drinking water is replaced for 2 days without DSS. Mice were sacrificed on the 7^th^ day by cervical dislocation. Colonic length and macroscopic damage Wallace score were recorded. Body weight loss and disease activity index were monitored during the whole procedure(Carle et al., 2023).

#### Oxazolone-induced colitis

12-week-old mice were pre-sensitized with 200 μL of 2% (w/v) 4-ethoxymethylene-2-phenyl-2-oxazoline-5-one (oxazolone) in 100% ethanol by subcutaneous injection. Five days later, mice were instilled intra-rectally with 150μL of 1.5 % oxazolone dissolved in 50% ethanol. For control mice, oxazolone was replaced with saline solution. Body weight loss and disease activity index were monitored every day (Al Nabhani et al., 2019).

### Paracellular permeability measurement

To measure the intestinal permeability, biopsies from duodenal and ileal mucosa were mounted in a Ussing chamber exposing 0.196 cm^2^ of tissue surface to 1.5ml of circulating oxygenated Ringer solution at 37°C. Paracellular permeability was assessed by measuring the mucosal-to-serosal flux of 4 kDa FITC-dextran (Sigma, France) (Al Nabhani et al., 2017b; Jung et al., 2012).

*In vivo* permeability assays were performed using fluorescein isothiocyanate (FITC)– dextran 4kDa as a paracellular permeability tracer. FITC–dextran was given by gavage to adult mice 6 days after initiation of the second cycle of DSS in the DSS-induced colitis model. Mice were gavaged with FITC–dextran (5 mg/200 μL/mouse) 4 h prior to sacrifice. Whole blood FITC–dextran concentration was determined by a spectrometry. FITC–dextran concentrations in serum were calculated from standard curves generated by serial dilution of FITC–dextran. *In vivo* permeability was expressed as the mean whole blood FITC–dextran concentration in ng/mL.

### ELISA

Biopsies of duodenum, ileum and colon from different mice models were collected and washed with cold PBS. These biopsies were then homogenized using an ultra-thurax in 1 ml of PBS1X and the concentration of proteins was determined using commercial kit (Biorad, Marnes la Coquette, France). IFN-γ, IL4, Il5, IL-1β, IL-12, IL13 and TNF-α protein levels in the intestine were determined by ELISA according to manufacturer’s instructions (BD Biosciences) (Meinzer et al., 2012).

### RNA extraction and real time quantitative PCR

After extraction by the NucleoSpin RNA II Kit (Macherey-Nagel, France), total RNAs were converted to cDNA using random hexonucleotides and then used for RT-PCR (Invitrogen, France). We conducted qPCR with QuantiTect SYBR Green PCR Kit (Applied, France) using sense and antisense primers specific for G3PDH, the long MLCK isoform (specifically expressed by epithelial cells), *Ifng*, *Il1b*, *Il4*, *Il5*, *Il12*, *Il13*, *Nod2*, *Mylk*, *Tnfa*, *Tnfr1*, *Tnfr2*, *Tjp1* and *Tjp2* (primers used available in table 1). The cycle threshold (Ct) was defined as the number of cycles at which the normalized fluorescent intensity passed the level of 10 times the standard deviations of the baseline emission calculated on the first 10 PCR cycles. Results are expressed as 2^−ΔΔCt^ as previously described (Al Nabhani et al., 2016).

**Table 1.**
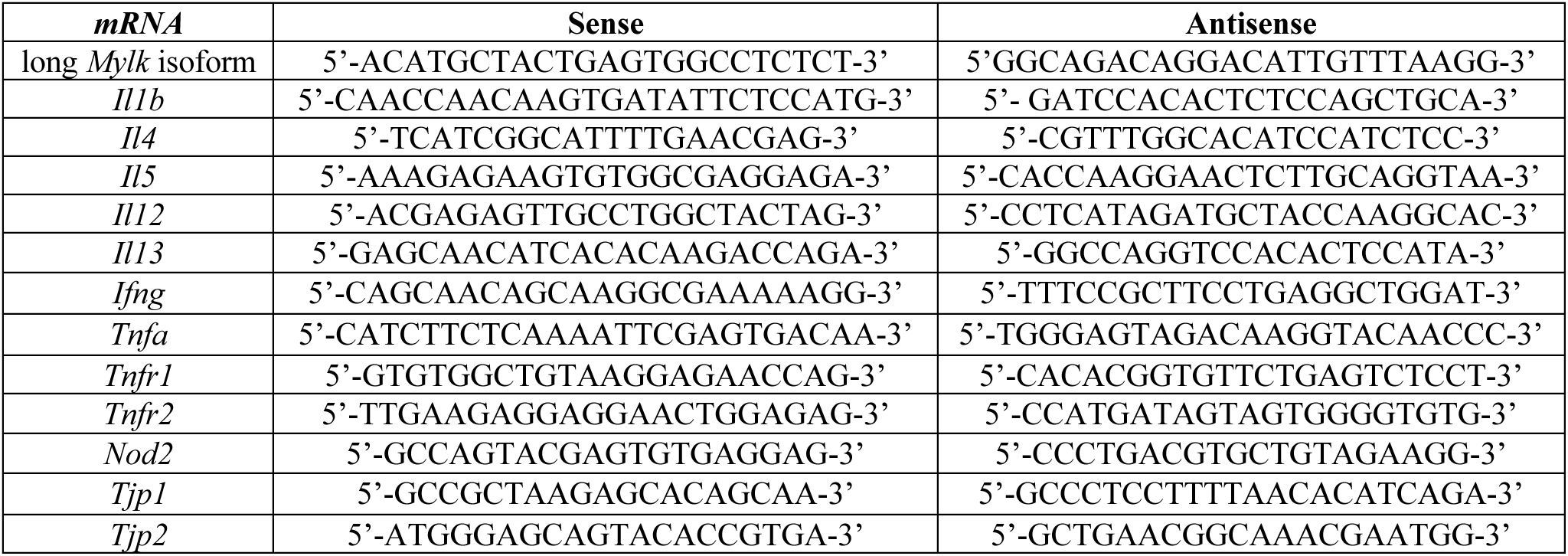
List of primers used for qPCR analyses in mice.

### Flow Cytometry

Small intestinal and colonic lamina propria cells were isolated as described previously (Al Nabhani et al., 2019). Briefly, Peyer’s patches were removed, and whole small intestine and colon were opened longitudinally, cut into pieces and incubated for 4 times for 8 min in EDTA (5mM), washed extensively, incubated for 20-30 min with 0.5mg/ml collagenase VIII (Sigma: c2139) and 10 U/ml DNase I (Roche 04536282001) and passed through a 100µm filter to generate single cell suspensions. Cells were separated by a 40/80% (w/v) Percoll density gradient and washed prior to staining. Cells were stained 10 min at 4°C with fixable Viability Dye eF450 (eBioscience Cat. No. 65-0863-14), to differentiate live cells from dead cells. Cells were further pre-incubated with Fc-Block for 10 min and stained for 20 min with antibodies to surface markers. For intracellular staining, cells were fixed and permeabilized with eBioscience Foxp3/Transcription Factor Staining Buffer Set (Invitrogen Cat. No.: 00-5523-00). Intracellular staining was performed with Alexa Fluor (AF)700-conjugated Foxp3 (clone FJK-16s, ThermoFisher) or with AF488-conjugated Gata3 (clone TWAJ, ThermoFisher) and AF647-conjugated RORγt (clone Q31-378, BD).

### Histological analysis

To perform histological analysis, small intestine rolls were established. The small intestine was enrolled, fixed in formol for 48 hours, then washed in 70% ethanol at room temperature. After inclusion in paraffin, sections of 5 µm were made for hematoxylin/eosin-Orange G coloration. Photos were acquired with Panoramic Scan Flash III and analyzed with CaseViewer. The length of at least 50 villi by sample, from the entire sample, was measured, and their goblet and immune cell number were counted. The histological analysis was also accompanied by determining the number of blunted villi per sample.

### Statistical analysis

For all the analysis, the Gaussian distribution was tested by the Kolmogorov-Smirnov test. Multigroup comparisons were performed using one-way ANOVA statistics with Bonferroni correction for multiple comparisons. For 2-group comparisons, an unpaired t-test assuming the Gaussian distribution was applied. Statistical analyses were performed using GraphPad Prism 7.00 (GraphPad Software). A two-sided P-value < 0.05 was considered statistically significant. All authors reviewed the data and approved the final manuscript.

## Abbreviations

CD: Crohn’s disease
DSS: Dextran Sulfate Sodium
MDP: Muramyl dipeptide
MLCK: myosin light chain kinase
NOD2: nucleotide oligomerization domain 2
Oxa: Oxazolone
TNBS: 2,4,6-Trinitrobenzenesulfonic acid
TNF-R: TNF receptor;
WT: wild-type.

## Acknowledgements

Z.A.N received funding from the European Research Council Starting Grant [WePredict project number: 949613], the Kenneth Rainin Foundation, the Helmut & Horten Foundation [Project ID: 2021-YIG-083], the Swiss National Science Foundation [SNSF, grant number: 310030_215675], the Krebsforschung Schweiz [KFS-5691-08-2022], the Jubiläumsstiftung von Swiss Life, the Sassella Stiftung, the Ruth & Arthur Scherbarth Stiftung, and the Novartis Foundation. We also thank Inselspital for technical and financial support. E.M. received funding from the French DGOS for the Rare Center of Rare Digestive Disorders. F.B. supported by INSERM, Université de Toulouse and the Association François Aupetit. J.R.T supported by NIH (grant number: R01DK068271). We also thank the Experimental Histopathology Facility of the INSERM / UPS / ENVT US006 CREFRE-Anexplo, Toulouse Purpan, France for technical assistance.

## Author contributions

Study design and concept: F.B., J.P.H., Z.A.N.; Data acquisition: A.M., F.B., F.C., G.D., G.S., K.B., L.M., M.R., N.N., R.A.A., T.N., Z.A.N.; Analysis and interpretation: A.J.M., A.M., E.M., F.B., F.C., J.P.H., J.R.T., L.M., M.R., T.N., Z.A.N.; Writing of the manuscript: A.M., T.N., F.B., Z.A.N.; Obtained funding: E.M., F.B., J.R.T., Z.A.N.; Technical support: A.M., F.C., G.D., G.S., K.B., L.M., N.N., RAA, TN; Study supervision: F.B., Z.A.N.

## Authors have no conflict of interest to declare

