## Supplementary Figures for "Colitis-primed circulating T cells drive small intestinal barrier function through Nod2, microbiota and myosin light chain kinase-dependent mechanisms"

#### Supplementary Figure 1

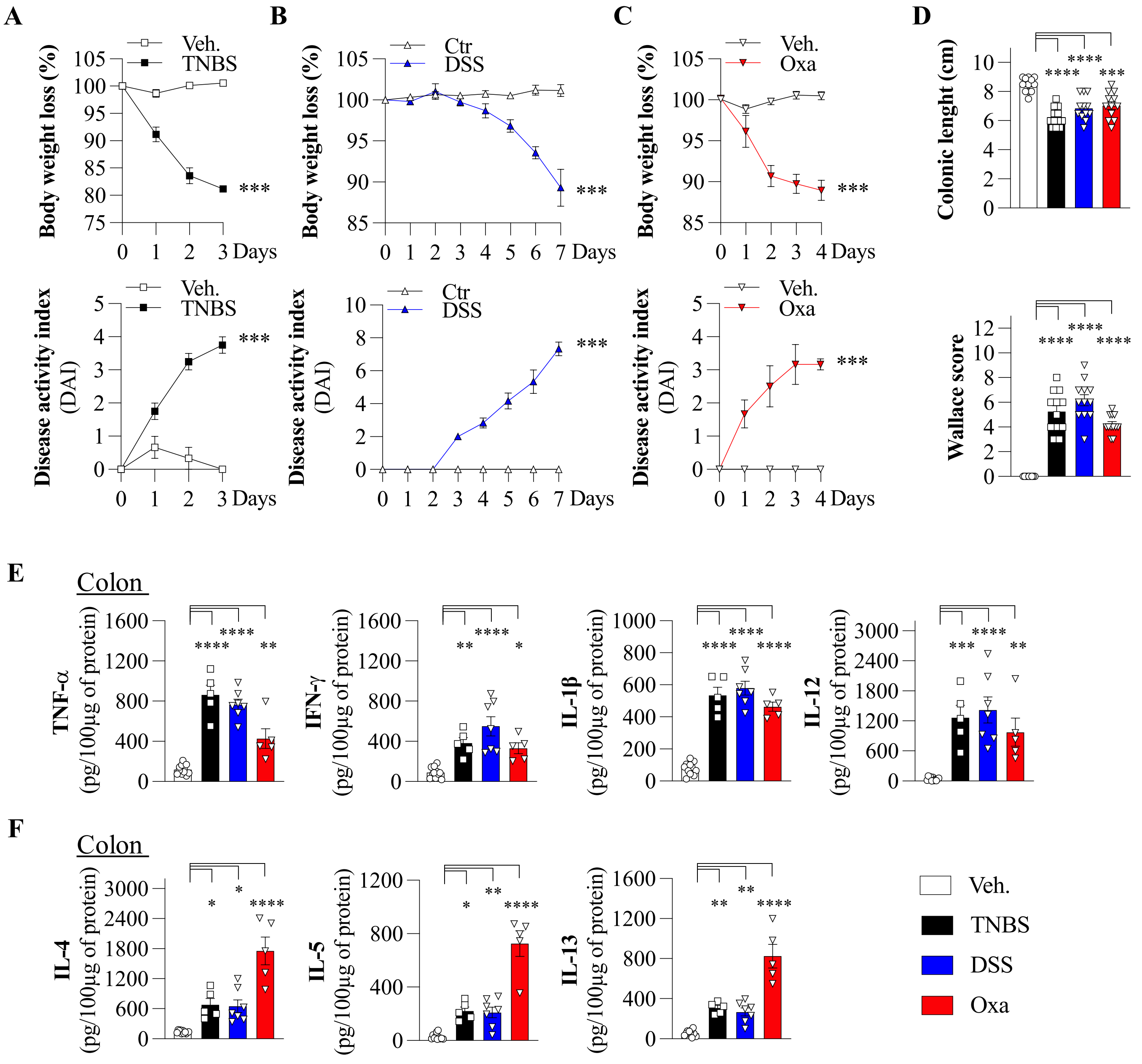

#### Supplementary Figure 2

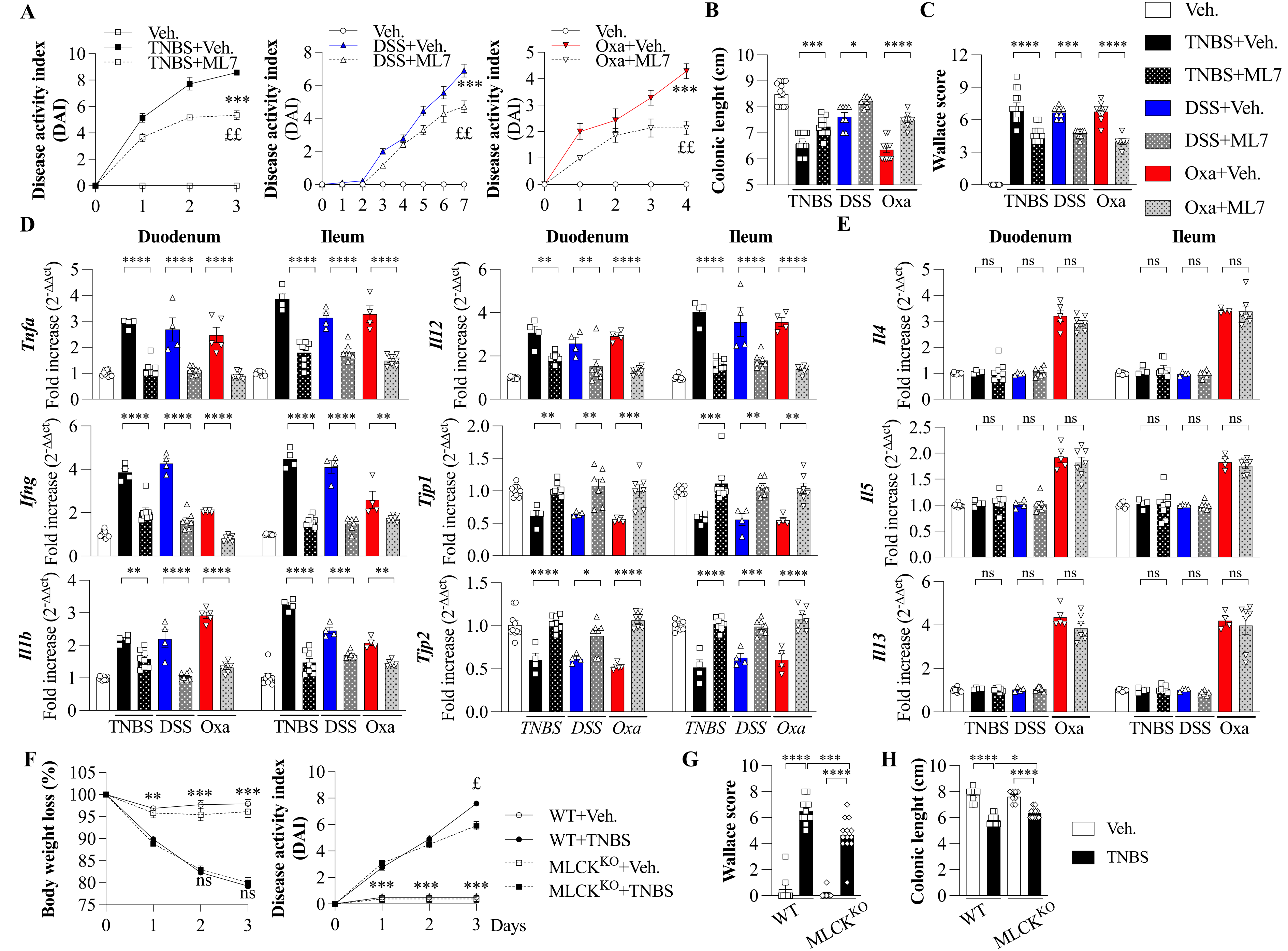

Supplementary Figure 3

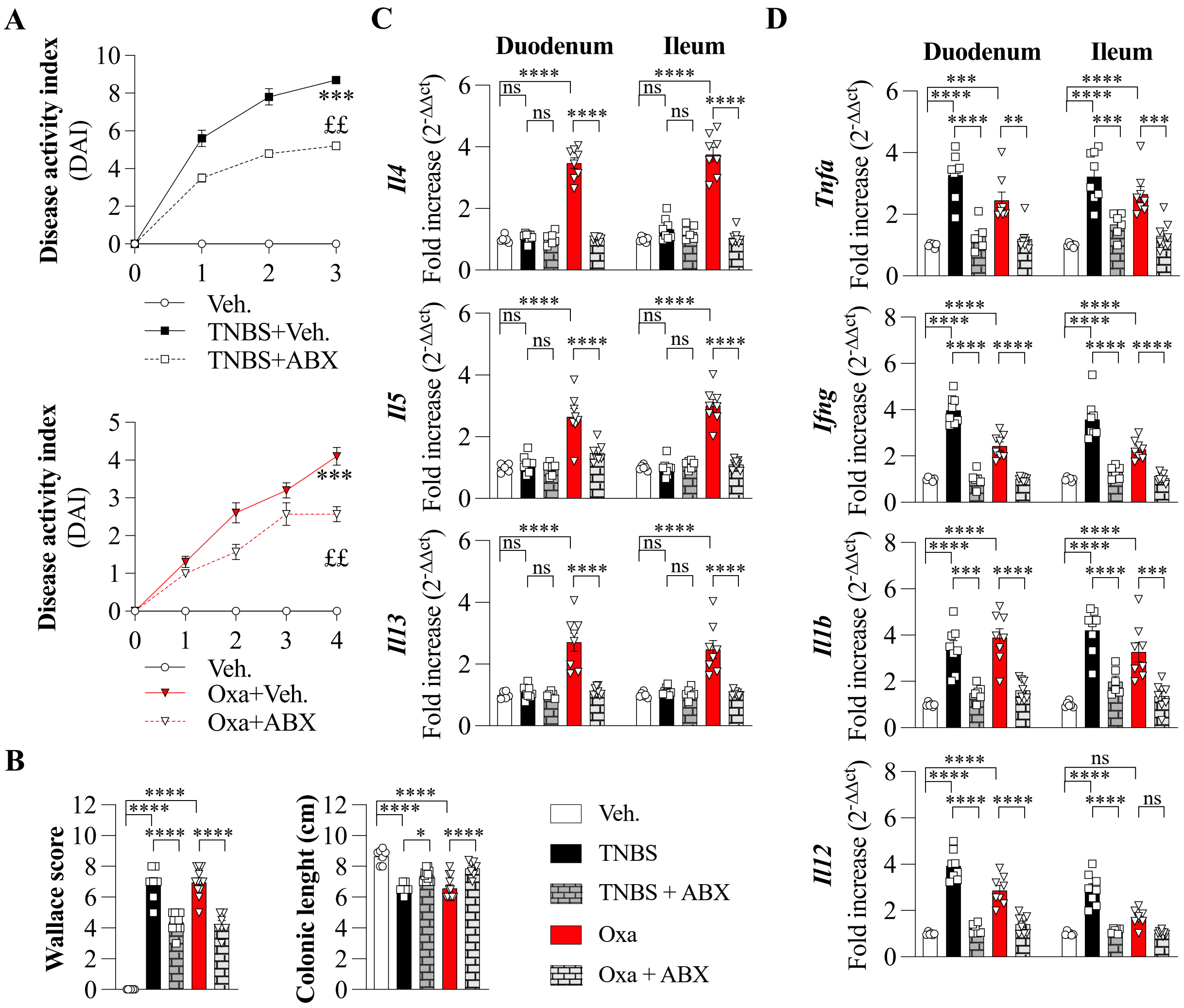

**Supplementary Figure 4**

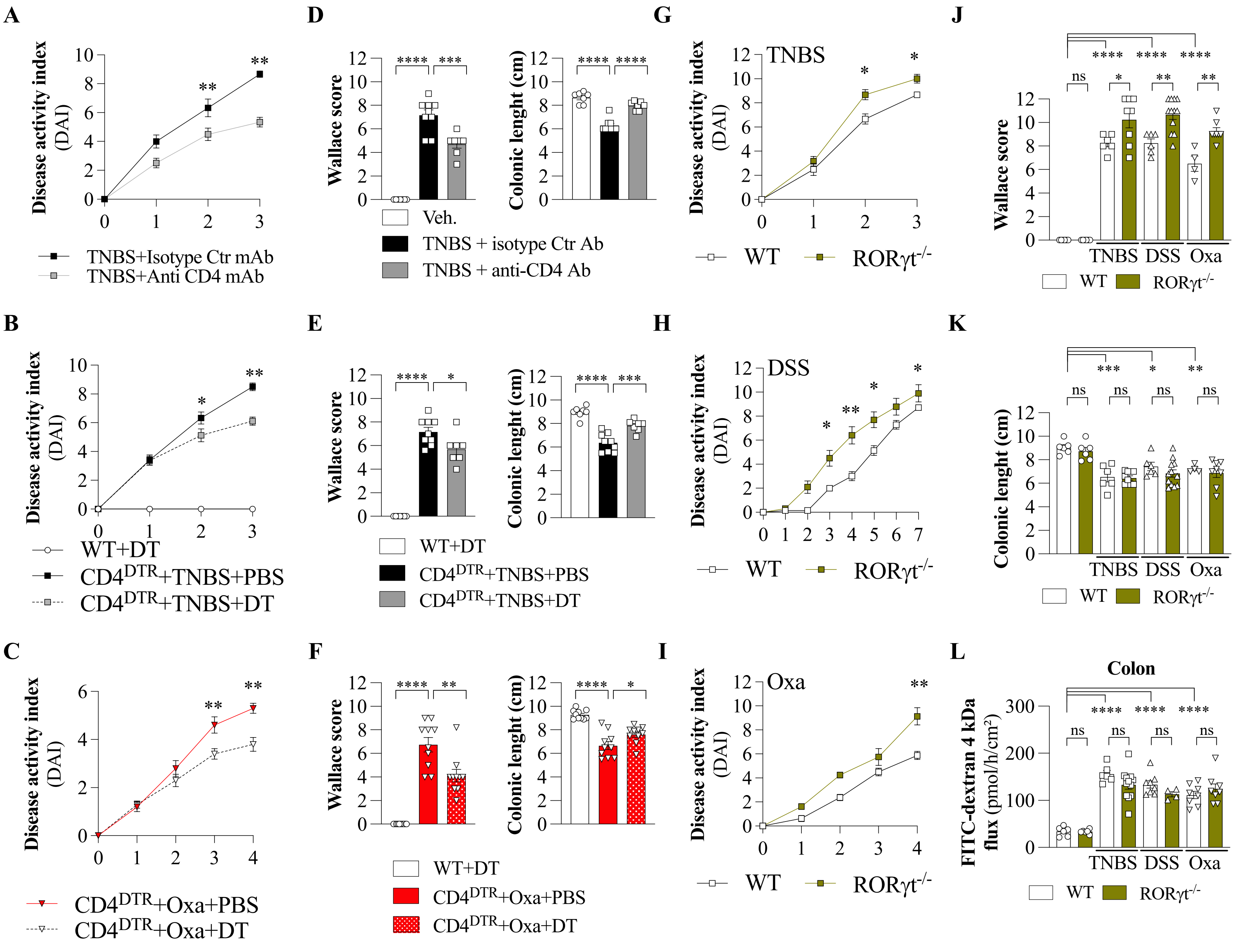

**Supplementary Figure 5**

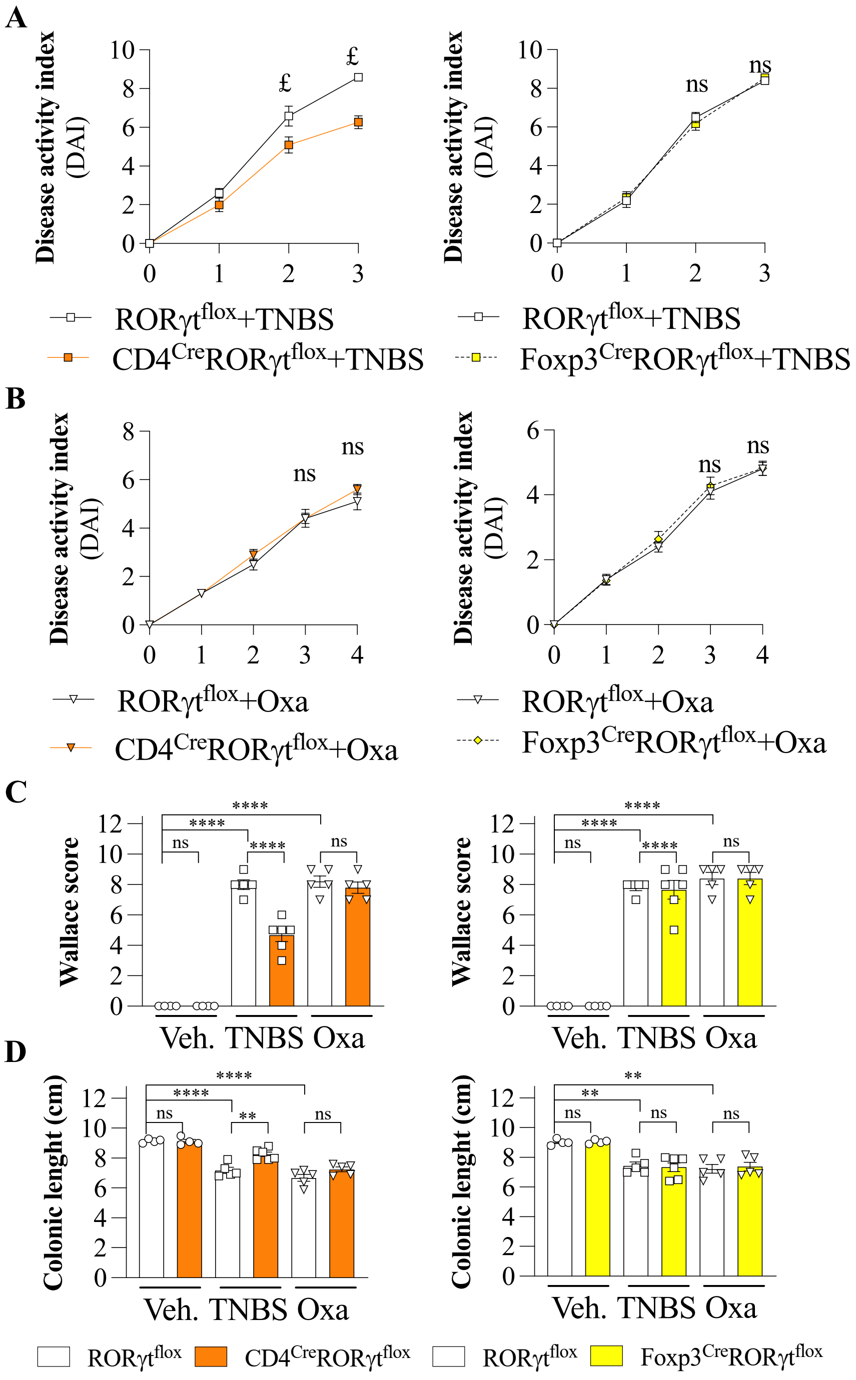

**Supplementary Figure 6**

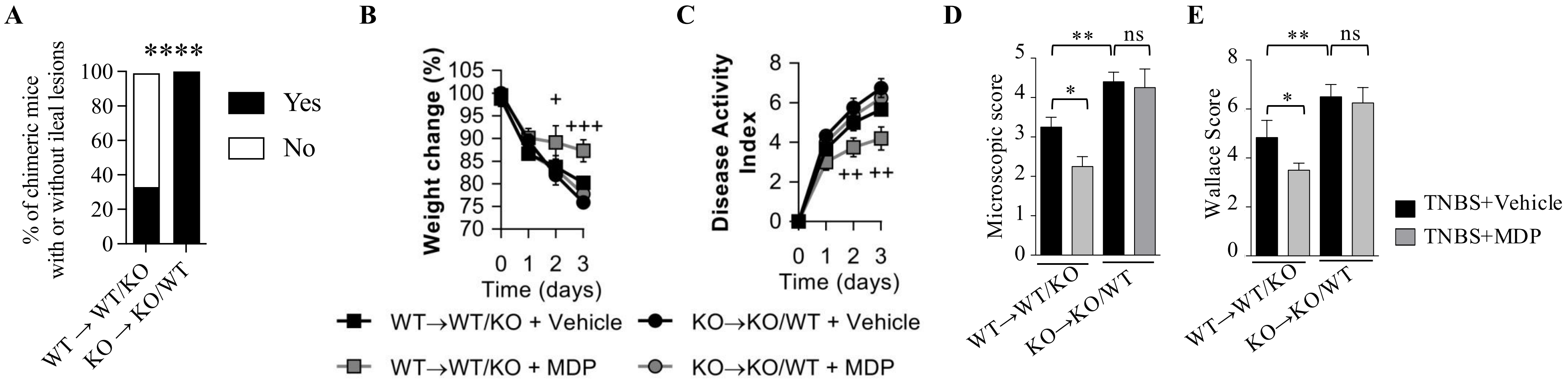

### Supplementary Figure 7

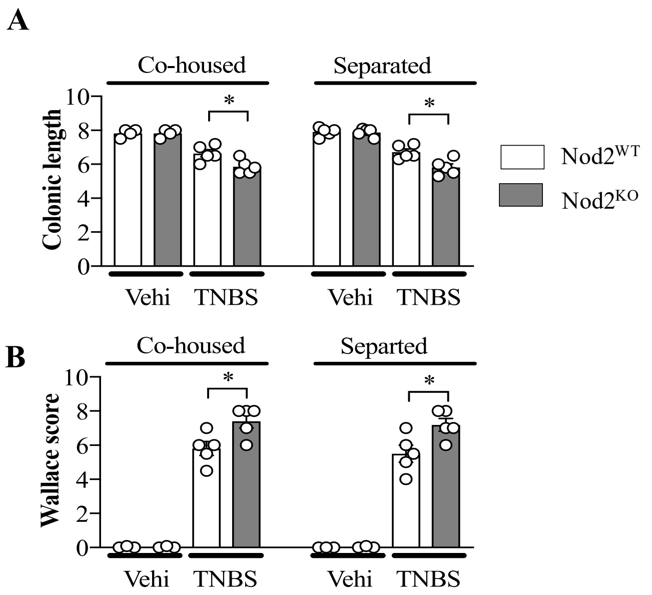

**Graphical Abstract**

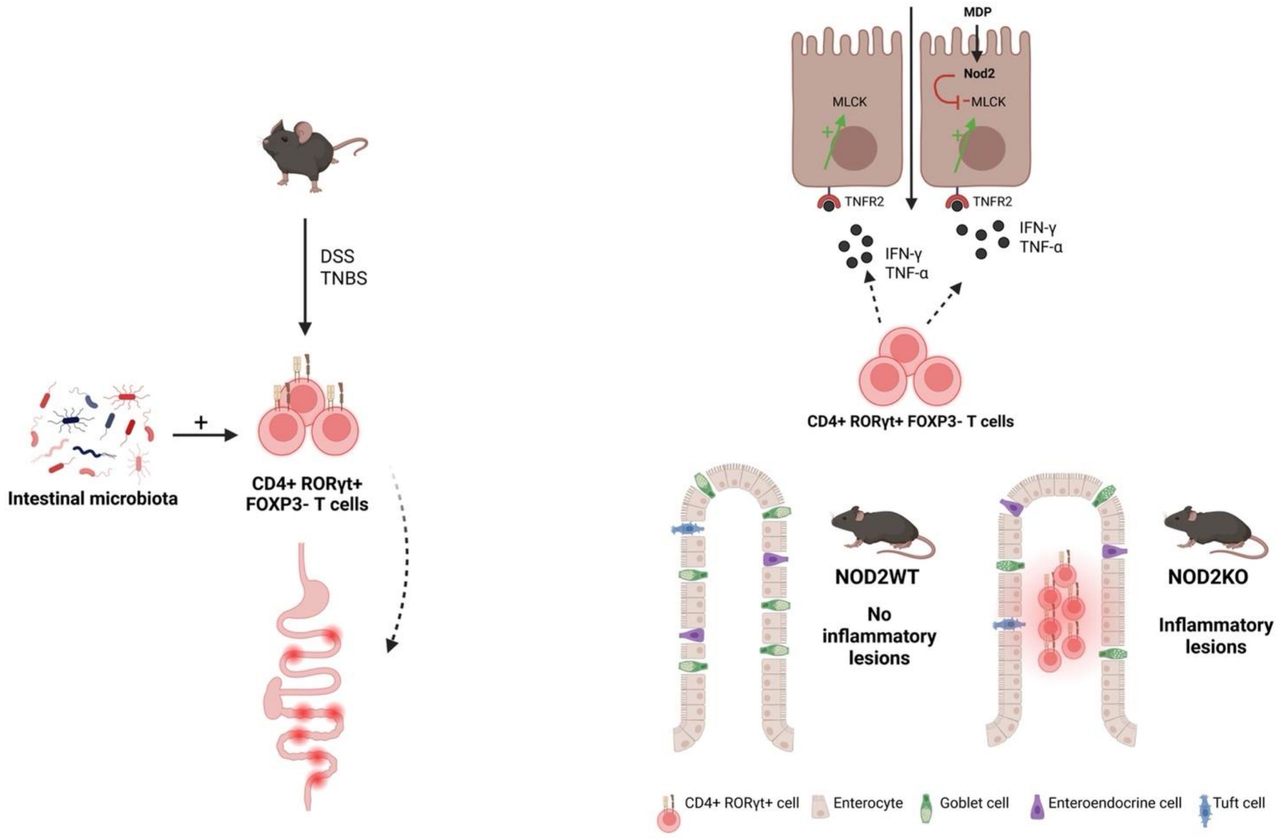
